# A Step-wise, Deterministic and Fatal Mouse Model of Myeloid Neoplasm with Spontaneous Acquisition of Patient-relevant RTK–RAS Mutations

**DOI:** 10.1101/2025.07.18.665518

**Authors:** Marija A. Zarocsinceva, Moosa Qureshi, Nicola K. Wilson, George Giotopoulos, Katherine H. M. Sturgess, M. S. Vijayabaskar, Sarah J. Kinston, Ryan Asby, David J. Adams, Brian J.P. Huntly, Fernando J. Calero-Nieto, Berthold Gottgens

## Abstract

Leukaemia arises through the stepwise transformation of healthy haematopoietic cells, yet the asymptomatic premalignant phase and its progression to overt disease remain poorly understood. To model this process, we engineered a patient-derived *CEBPA* mutation into Hoxb8-FL multipotent murine progenitors and transplanted them into syngeneic mice, capturing a clinically silent premalignant stage. All recipients developed overt disease after ∼12 months with 100% penetrance and all acquired secondary RTK–RAS mutations, often with identical amino acid changes to those in patients. Single-cell transcriptomics and phenotypic profiling showed that premalignant mutant cells adopt a plasmacytoid dendritic progenitor–like state in vitro which generates both myeloid and B-lymphoid lineages during premalignancy in vivo, with individual tumours restricted to one lineage. Specificity for RTK-RAS mutations coupled with ongoing differentiation is reminiscent of Juvenile Myelomonocytic Leukaemia, where cellular models are currently lacking, thus providing a tractable model of myeloid neoplasm for mechanistic studies and drug discovery.

## Introduction

Cancer evolution is a dynamic process often spanning many years before it is detected in the clinic. Advanced genomics technologies have been able to reconstruct this process with great precision in the context of leukaemia, where extrapolation from single cell derived genome sequencing data has provided strong evidence that initial preleukaemic mutations for both paediatric and adult leukaemia may already have occurred *in utero* ^1,2^. Leukaemogenesis is therefore commonly a slow process, whereby stepwise acquisition of mutations provides increased fitness to individual preleukaemic clones, ultimately leading to the development of aggressive disease.

While genomics can reconstruct stepwise trajectories of tumour evolution retrospectively, the early stages of leukaemogenesis are typically inaccessible in the human setting where individuals are asymptomatic. Importantly, a better understanding of the preleukaemic phase will have important clinical implications. Deep sequencing of acute leukaemia samples has revealed that genetically diverse blast populations often derive from a single founding clone ^3–5^ and the same preleukaemic mutations that were present in the initial tumour are commonly found in relapse samples ^5–7^. This suggests that preleukaemic cells can evade cytotoxic chemotherapy yet retain the capacity to regenerate the disease. Although often present only in very low numbers and typically outnumbered by highly proliferative leukaemic blasts, these cells often go undetected in bulk tumour analyses. New experimental models to study cancer progression from its earliest stages are therefore essential to gain a comprehensive understanding of both initial tumour evolution, as well as to design interventions that target the disease at its root, thereby improving patient outcomes.

Retroviral transduction models have long been a mainstay of leukaemia modelling ^8,9^, whereby a pool of mouse bone marrow stem/progenitor cells are transduced with a retrovirus carrying a leukaemia oncogene *ex vivo*, followed by transplantation to track disease development in an *in vivo* setting. One potential drawback of this approach is the substantial heterogeneity of cell states at the time of transduction, essentially obscuring the precise identity of the initiating cell. To better control for the nature of the initiating cell state, we have previously taken advantage of conditionally immortalised Hoxb8-FL cells ^10^ which represent a functional *in vitro* counterpart of lymphoid-primed multipotent progenitors (LMPP) that can be maintained in culture by combining the known self-renewal properties of *Hox* genes with the pro-survival function of the Flt3 ligand. Of note, LMPP have been suggested as one possible candidate cell of origin for AML as well as a potential normal counterpart to LSCs ^11^ thus underscoring the potential relevance of Hoxb8-FL cells for disease modelling. Using the strong *MLL-ENL* leukaemia oncogene, we therefore previously demonstrated how Hoxb8-FL cells provide a powerful cell system for *in vivo* leukaemia modelling, taking advantage of its homogeneity coupled with retained multilineage differentiation capacity ^12^.

The *CEBPA* gene is commonly somatically mutated in Acute Myeloid Leukaemia (AML) patients, with mutation of *CEBPA* believed to be an early leukaemogenic event ^13^. Moreover, relapse samples from AML patients usually retain the *CEBPA* mutation from the initial tumour ^14^ and germline *CEBPA* variants may also affect development of AML, highlighting early and disease initiating roles for CEBPA. An earlier retroviral transplantation model testing a range of *CEBPA* mutations reported that a *CEBPA*-N321D mutant caused the most rapid and aggressive disease ^15^. Here we combined the Hoxb8-FL cellular model with a *CEBPA*-N321D retrovirus to capture the effects of *CEBPA* mutation on the haematopoietic system in a tightly controlled setting, and thus provide new insights into early cellular and molecular steps of leukaemia development.

Our new tractable model of leukaemia development shows how *CEBPA* N321D can substitute for the self-renewal function of Hoxb8 *in vitro* and generate a premalignant state when transplanted *in vivo*. Progression to an aggressive disease *in vivo* entails a long latency period and requires the acquisition of secondary driver mutations, exquisitely targeting five members of RTK-RAS pathway. The new model thus captures all critical stages of leukaemogenesis, for which we report comprehensive mutation-, as well as single cell gene expression states. Our new model provides a utilitarian tool, by combining a well-defined cell of origin with deterministic evolution from preleukemic *CEBPA* N321D cells to fully transformed leukaemic cells harbouring clinically relevant and somatically acquired RTK-RAS pathway mutations, all accessible for mechanistic and therapeutic experimentation in a syngeneic fully immunocompetent *in vivo* setting.

## Results

### N321D mutant *CEBPA* endows Hoxb8-FL cells with Hox-independent propagation

A common drawback of experimental model systems to study leukaemia initiation and progression is the phenotypic and functional heterogeneity of the starting population. Hoxb8-FL cells on the other hand provide a relatively homogeneous pool of progenitors. Based on the recognised significance of *CEBPA* C-terminal bZIP mutations in leukaemia, we therefore created a cellular model with the *CEBPA* N321D mutation in Hoxb8-FL cells. To this end, multipotent Hoxb8-FL cells were transduced with retroviral MSCV-GFP constructs carrying either the mutant or wild type *CEBPA* (*CEBPA* N321D and *CEBPA* WT respectively), or an empty vector control (EV) (**Figure 1A**). To allow time for retroviral integration into the Hoxb8-FL host genome, cells were expanded for two days in self-renewal conditions (β-oestradiol to ensure expression of *Hoxb8* and Flt3L) but subsequently transferred to differentiation conditions lacking oestrogen (hereafter Flt3L). Only the *CEBPA* N321D mutant cells continued to proliferate throughout the duration of the experiment in Flt3L media demonstrating that expression of the mutant *CEBPA* N321D conferred *Hox*-independent self-renewal capacity (**Figure 1B**). Our results with EV and *CEBPA* WT cells by contrast are consistent with the original Hoxb8-FL publication ^10^ where the authors showed that following β-oestradiol withdrawal, cells expanded for 6 days, differentiated, and remained viable for only 9 days.

**Figure 1.**
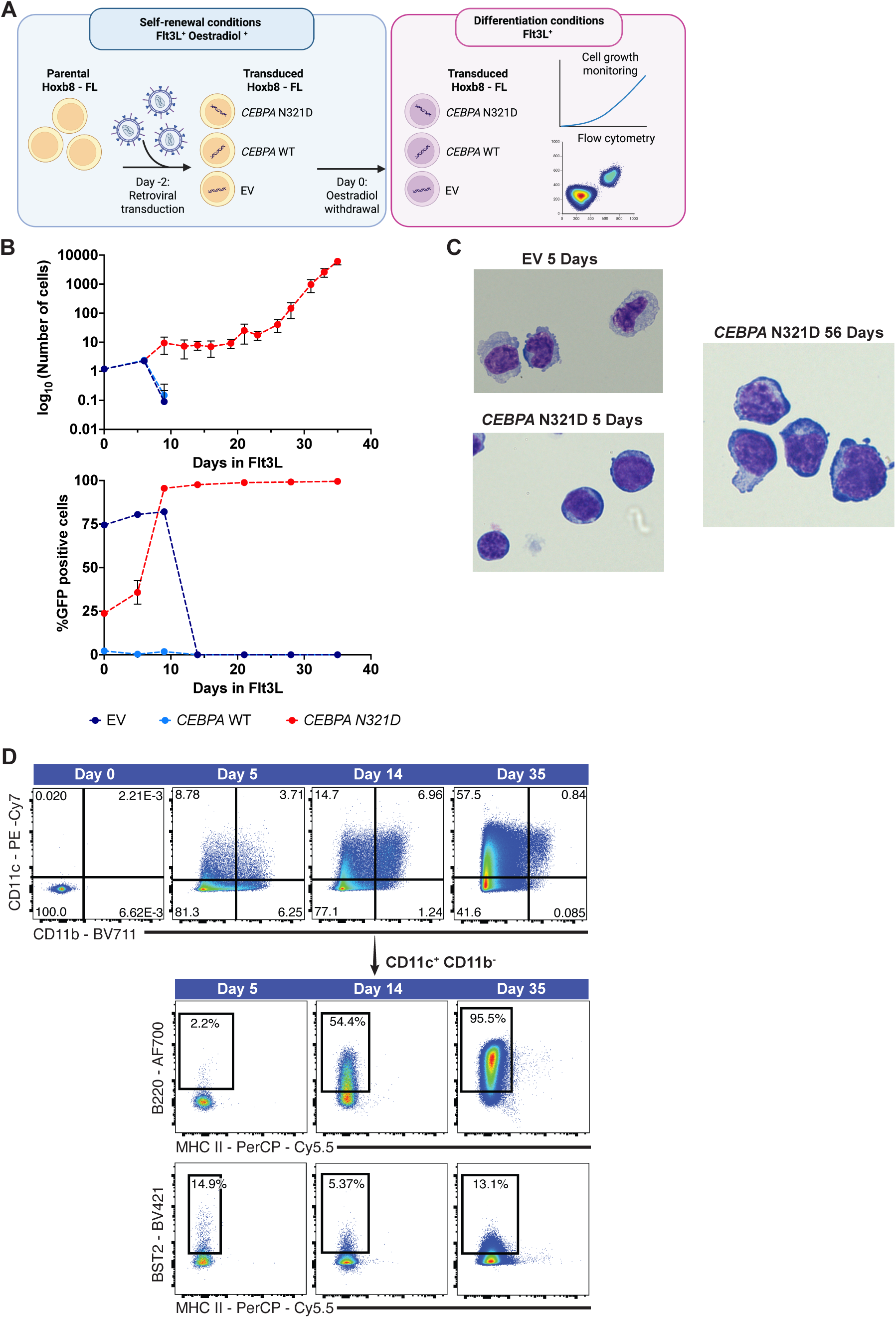
N321D mutant *CEBPA* endows Hoxb8-FL cells with Hox independent propagation. **A.** Schematic of establishment and characterisation of *CEBPA* N321D model. Hoxb8-FL cells were transduced with different retroviral constructs and subjected to β-oestradiol withdrawal (abrogating Hoxb8 signalling). Differentiation potential of the resulting cells was assessed by monitoring cell growth and characterising haematopoietic marker expression via flow cytometry. **B.** Cell numbers and the percentage of GFP-positive cells over time for transduced cells cultured in Flt3L (differentiation conditions). The displayed values were calculated as the mean of n=2 for EV and *CEBPA* WT and n=3 for *CEBPA* N321D. The error bars are standard deviation. **C.** Brightfield images of Romanowsky stained GFP^+^ EV and GFP^+^ *CEBPA* N321D cells cultured in Flt3L media for 5 days and of *CEBPA* N321D cells cultured in Flt3L media for 56 days. **D.** Flow cytometry analysis of dendritic cell marker expression in *CEBPA* N321D cells in Flt3L media.

**Supplementary Figure 1.**
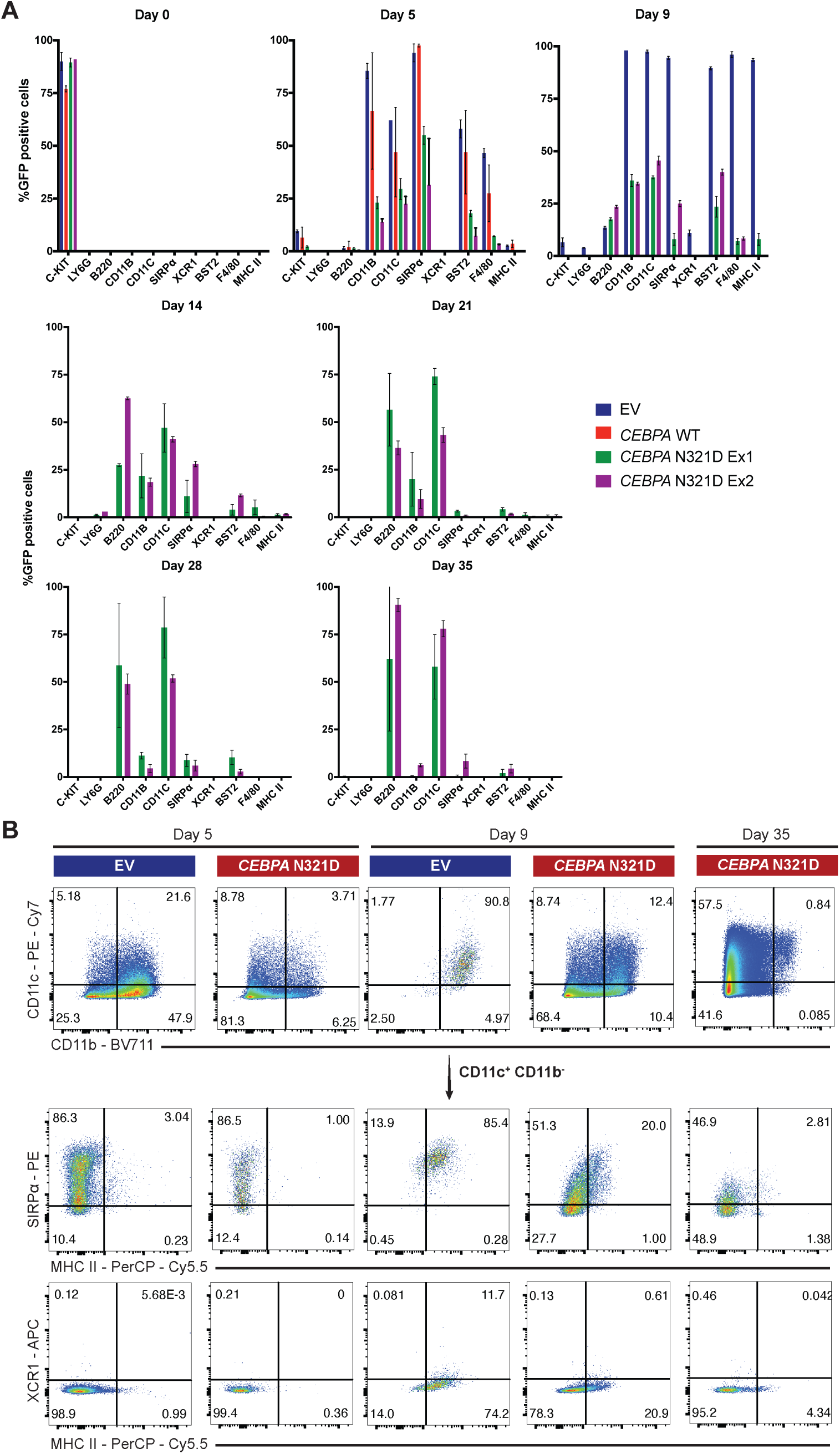
**A.** Expression of all sampled haematopoietic markers in *CEBPA* WT, EV and *CEBPA* N321D cells. Two independent experiments of *CEBPA* N321D cells generation are included (Ex1 and Ex2 respectively). Each bar plot shows the calculated mean of two technical replicates and error bars represent the standard deviation. EV, *CEBPA* WT and *CEBPA* N321D Ex1 were performed in the same experiment and *CEBPA* N321D Ex2 repeated in a separate experiment. **B.** Expression of cDC haematopoietic markers in EV and *CEBPA* N321D transduced Hoxb8-FL cells cultured in Flt3L media over time (days). A small number of *CEBPA* N321D cells expressed both CD11c and CD11b at day 5 and day 9 in Flt3L conditions but this population was almost absent at day 35. This contrasts with the EV cells that showed a much higher CD11c^+^ CD11b^+^ population at day 5 and day 9. Interestingly, EV cells gave rise to the SIRPa^+^ cDC2 subtype but were unable to differentiate into the XCR1^+^ cDC1 subtype. There was only a very small number of SIRPa^+^ MHC II^+^ cells in the *CEBPA* N321D CD11c^+^ CD11b^+^ population present in long-term culture.

**Supplementary Figure 2.**
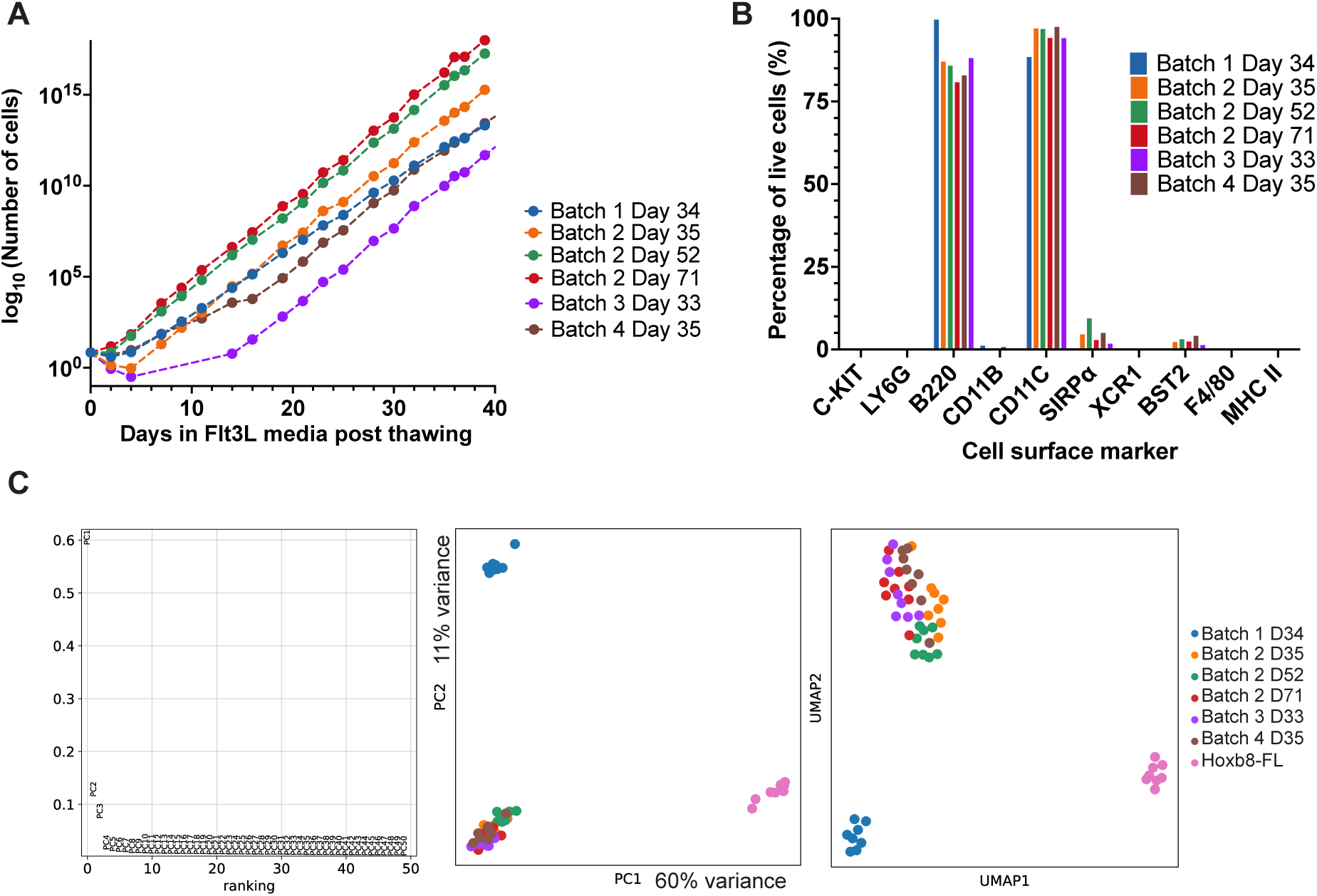
**A.** Growth curves of independent replicates of *CEBPA* N321D cells grown in Flt3L media. **B.** Expression of haematopoietic markers as determined by flow cytometry analysis. Cells were collected after 38 days in Flt3L media post thawing. All samples are B220^+^ CD11C^+^. A very small number of cells also showed expression of SIRPα and BST2 markers, apart from cells from Batch 1 day 34. **C.** Left: variance explained by each of the calculated PCs; middle: PCA plot showing PC1 and PC2; right: UMAP representation of the data coloured by batch. D - indicates number of days cultured in Flt3L media post thawing.

### Hox-independent mutant CEBPA cells adopt a dendritic progenitor phenotype

Morphological examination at day 5 post β-oestradiol withdrawal demonstrated that EV cells displayed classic dendritic cell features including lateralised reniform nuclei, vacuolised cytoplasm and a few dendritic projections (**Figure 1C**). By contrast, these features were absent in the *CEBPA* N321D cells analysed at the same time point. Of note, after 56 days in Flt3L conditions, a subset of *CEBPA* N321D cells exhibited some dendritic cell features (reduced cytoplasm:nuclei ratio, occasional reniform nuclei and emergence of some small dendritic protrusions), but they never displayed a fully mature dendritic cell morphology.

Redecke *et al.*^10^ showed that Hoxb8-FL cells grown in the presence of Flt3L differentiate into a mixed population of cDCs (defined by CD11c^+^, CD11b^+^, MHC II^+^, B220^-^) and pDCs (defined by CD11c^+^, CD11b^-^, B220^+^). Our *CEBPA* N321D cells showed an increase in the CD11c^+^ CD11b^-^ population over time (**Figure 1D**). This population also became progressively more positive for the pDC marker B220. Interestingly, the *CEBPA* N321D cells remained MHC II^-^ throughout the experiment and only lowly expressed BST2, suggesting that they remain in a stalled progenitor-like state (**Supplementary Figure 1A, 1B)**. In addition, we performed low-cell number RNA-sequencing, sampling four batches of *CEBPA* N321D cells at sequential time points during Flt3L culture. The four batches, produced in independent experiments, showed a high degree of similarity in growth rates, levels of CD11c and B220 expression as well as transcriptomic homogeneity, thereby confirming high reproducibility of generating the Hox-independent *CEBPA* N321D self-renewing phenotype (**Supplementary Figure 2A-2C**). Collectively, these results demonstrate that *CEBPA* N321D cells have a defect in DC differentiation and instead acquire a self-renewing pDC progenitor-like phenotype.

### *CEBPA* N321D cells give rise to a long-latency myeloid neoplasm *in vivo*

To validate if *CEBPA* N321D cells had leukaemogenic potential *in vivo*, we transplanted GFP+ *CEBPA* N321D or EV cells into lethally irradiated recipients (**Figure 2A**). When peripheral blood was drawn from recipients 7 days post-transplantation (**Figure 2B**), GFP^+^ cells were detected in all recipients, albeit that higher chimerism was observed for *CEBPA* mutant cells (>10%) compared to the EV cells (<1%). No EV cells were detected past day 7, whereas *CEBPA* N321D cells were detected at all sampled time points, reiterating that EV cells are only capable of short-term repopulation, consistent with the original Hoxb8-FL study ^10^. The presence of a GFP positive population in the peripheral blood for such an extended period firstly demonstrated that the *CEBPA* N321D progenitor cell type displayed sustained self-renewal activity *in vivo*, and secondly produced output that could be detected in the peripheral blood. The sustained engraftment without the development of rapid disease onset also prompted us to hypothesise that secondary mutations were likely to be required for disease progression. To prospectively test this hypothesis, 4 of the 12 mice transplanted were subjected to an additional low dose (1Gy) of ionising irradiation (Irradiated, IR), as a further mutagenic stimulus, 3-4 months post the initial transplantation (**Figure 2A**).

**Figure 2.**
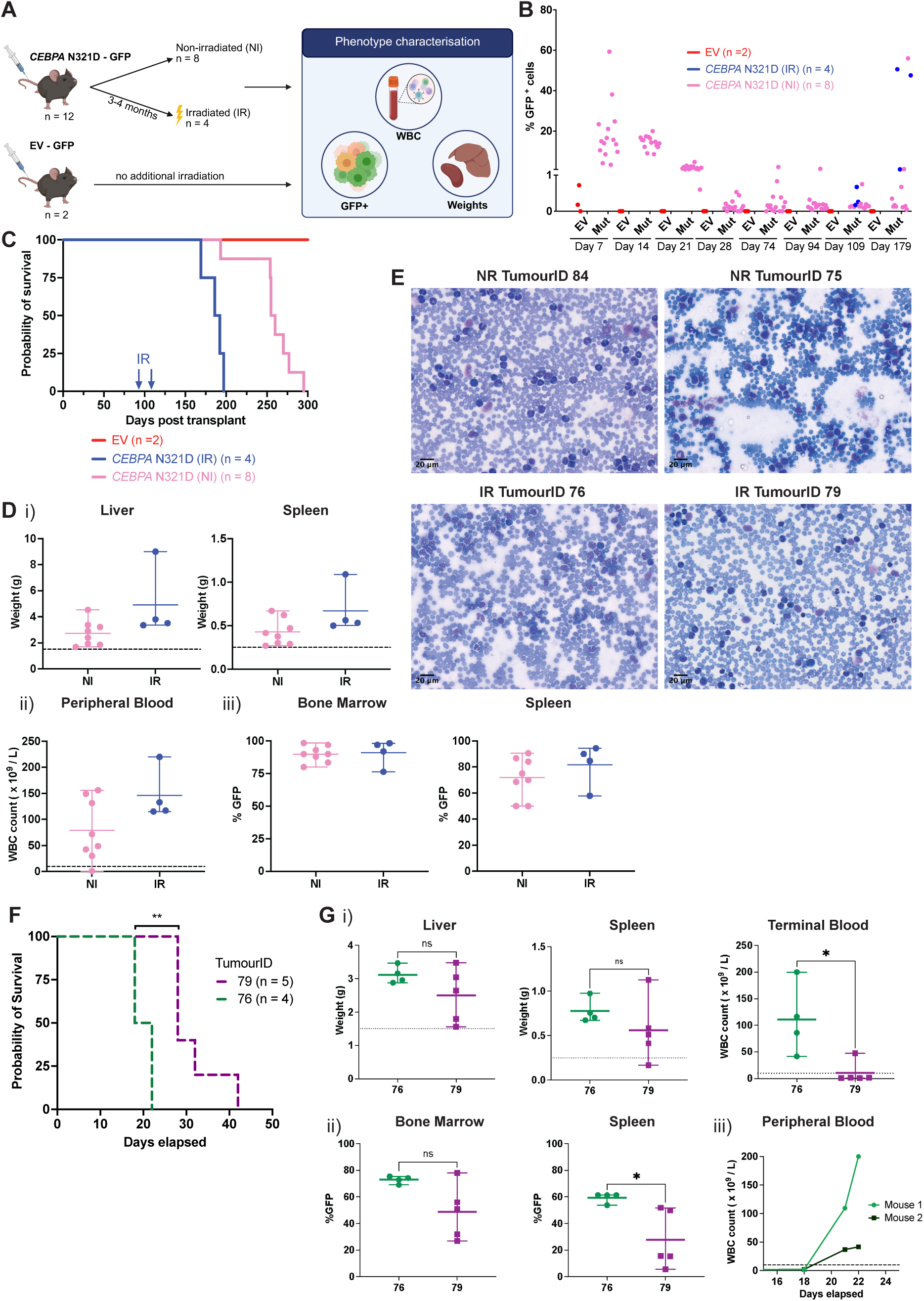
*CEBPA* N321D cells give rise to a long latency leukaemia *in vivo*. **A.** Schematic of mouse transplantation experiments to assess the leukaemogenic potential of the *CEBPA* N321D cells. Lethally irradiated mice were injected with either *CEBPA* N321D cells (n=12) or EV cells (n=2). Mice injected with *CEBPA* N321D cells were subdivided into two cohorts: Cohort non-irradiated, NI (n=8) received no additional doses of irradiation. Irradiated cohort, IR (n=4) received an additional low dose of (mutagenic) irradiation (1 Gy) 3 or 4 months after transplantation. Disease progression was assessed by measuring WBC and %GFP positive cells in the peripheral blood. Symptomatic mice were culled, and their liver and spleen were weighed. **B.** Peripheral blood chimerism of mice injected with *CEBPA* N321D or EV cells. **C.** Survival of primary recipients. **D.** Characterisation of the disease phenotype. i) Graphs show hepatomegaly and splenomegaly in symptomatic mice, ii) terminal WBC, iii) %GFP in BM and %GFP in spleen. **E.** H&E-stained peripheral blood smears from four symptomatic mice injected with *CEBPA* N321D cells. **F.** Survival curves of secondary recipients transplanted with BM cells from two primary tumours (Mantel-Cox test, **pvalue = 0.004). **G.** Characterisation of secondary disease phenotype. Unless otherwise stated, significance calculated with Mann Whitney test. i) Liver and spleen weights at time of disease presentation (ns p-values 0.413 and 0.190). Terminal WBC of symptomatic mice (*p-value = 0.032). ii) GFP% in bone marrow and spleen (p-values 0.190 and 0.016). iii) Time course of WBCs of two mice injected with cells from Tumour ID 76. Dashed lines in D) and G) indicate upper limits of normal liver weight (1.5g), spleen weight (0.25g) and WBC (x10^9^ / L).

**Supplementary Figure 3.**
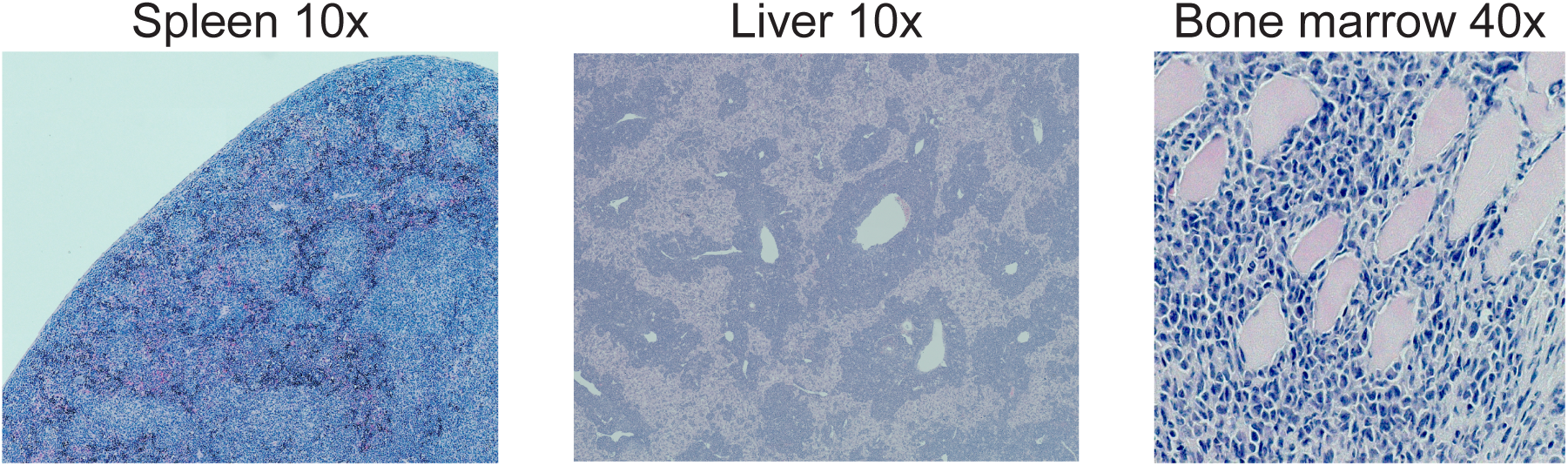
H&E-stained representative histology sections of liver, spleen and bone marrow from a mouse (Tumour ID 76) transplanted with the *CEBPA* N321D cells.

All mice injected with *CEBPA* N321D cells ultimately developed leukaemia, whereas no mice injected with control EV cells showed any signs of leukaemia development or sustained engraftment during the course of the study (**Figure 2C**). The non-irradiated (NI) cohort had a median survival of 257 days, whereas recipients from the IR cohort developed disease around 190 days post transplantation, in line with our prediction that IR would accelerate disease development. All symptomatic mice injected with *CEBPA* N321D mutant cells showed splenomegaly and hepatomegaly and elevated white blood counts (WBC), consistent with the phenotypic presentation of leukaemia (**Figures 2Di and 2Dii, Supplementary Figure 3**). Flow cytometry characterisation showed the presence of GFP positive cells in the bone marrow (>75%) and spleen (>50%) of all animals, confirming that the observed leukaemia was initiated by the transplanted mutant *CEBPA* N321D cells **(Figure 2Diii)**.

Peripheral blood smears from four representative mice demonstrated elevated numbers of cells with blast-like appearance (**Figure 2E**). To determine whether BM cells isolated from the diseased animals possess tumour-initiating capacity and self-renewal potential, we selected BM cells from two primary tumours (Tumour ID 76 and 79) and transplanted the cells into a new cohort of mice. All animals developed disease 18 to 42 days post-transplantation. Of note, a marked difference in disease latency was seen between the recipients of each tumour, potentially reflecting differences in their secondary driver mutations and/or leukaemic stem cell numbers (**Figure 2F**). Secondary recipients had increased liver and spleen weights, increased WBCs (all recipients of tumour 76, but not 79) (**Figure 2Gi**) and infiltration of GFP-positive cells in BM and spleen, consistent with a leukaemia phenotype (**Figure 2Gii)**. Highlighting the potential utility of our model for future mechanistic and therapeutic studies, we commonly observed an increase in the percentage of GFP^+^ cells as an early indicator of disease progression (**Figure 2Giii**).

### Leukaemia onset requires secondary mutations in the RTK-RAS pathway

The long latency of disease following transplantation of *CEBPA* N321D cells, and the observed acceleration of disease progression with mutagenic stress prompted us to test whether secondary mutations were required for progression into overt leukaemia. DNA was extracted from whole bone marrow of the 12 primary tumours, as well as the pre-injection CEBPA N321D cell line (as a reference) and subjected to mutational analysis by whole exome sequencing (WES) (**Figure 3A**). Included were also 2 tumour samples from a small pilot transplantation experiment that was performed following the same procedure. Exome sequencing of single nucleotide variants (SNVs) and small indels, identified a median value of 28 mutations per sample, with the IR samples showing slightly elevated numbers of mutations, as expected. However, despite their elevated mutational burden no mutations were specific to the IR cohort (compared to the NI samples) (**Supplementary Figures 4A and 4B**), suggesting that the CEBPA N321D may exert a strong selective pressure on the secondary drivers and that the additional mutagenic stress did not result in an obviously qualitatively different disease.

**Figure 3.**
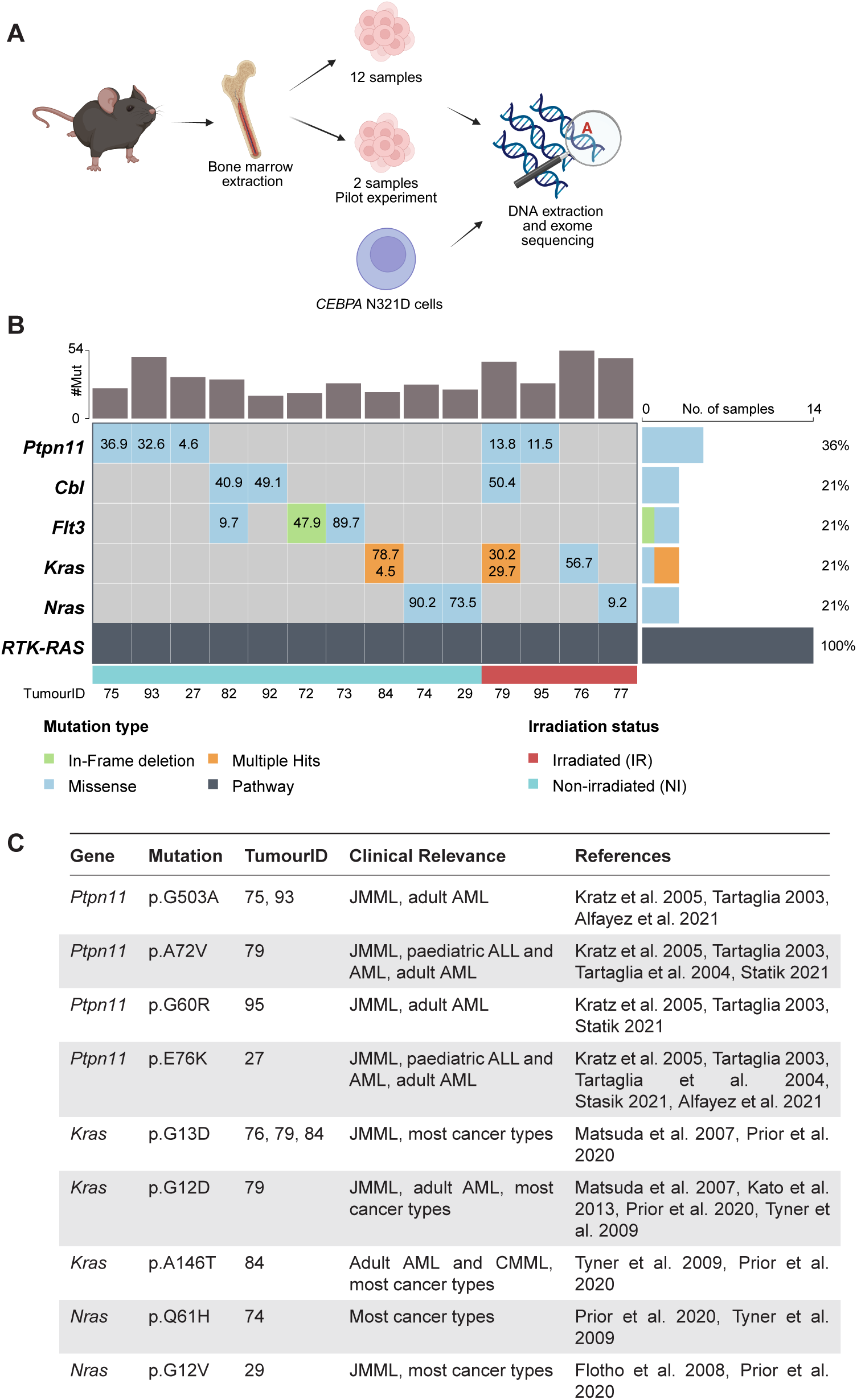
Leukaemia onset requires secondary mutations in the RTK-RAS pathway. **A.** Diagram showing the outline of the exome sequencing experiment. **B.** Mutations found in the genes of the RTK-RAS pathway. The bar chart on top of the plot shows the number of mutations found in each of the tumour samples. The bar chart on the right depicts the % of samples with mutations found in each of the 5 genes. The calculated VAF is displayed inside the box. If more than one mutation was found in a gene, the VAFs for each of the mutations are displayed together in the same box. **C.** Mutations found in this study that are frequently found in haematological malignancies ^17–22,24–27^.

**Supplementary Figure 4.**
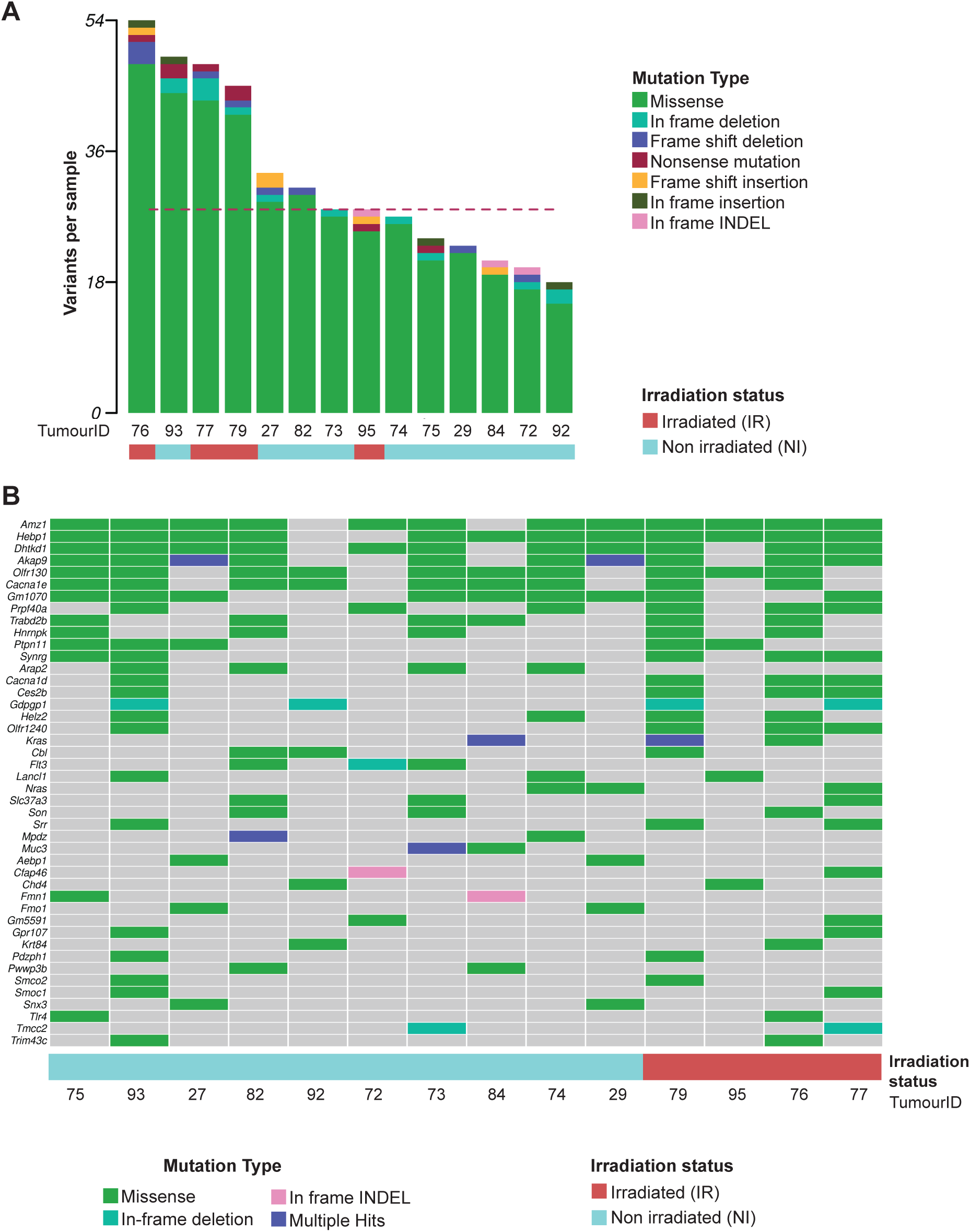
Mutations found in the *CEBPA* N321D BM cells. **A.** Bar chart showing the number and type of the acquired mutations in each of the 14 tumour samples. Irradiated samples Tumour ID 76, 77, 79 and 95 are marked in red. **B.** Oncoplot depicts mutations found in more than 2 samples. The presence of a mutation in a gene is shown by a non-grey colour, with the colour corresponding to a specific type of mutation. The genes are listed in order from highest to lowest based on the number of samples in which they are found to be mutated.

**Supplementary Figure 5.**
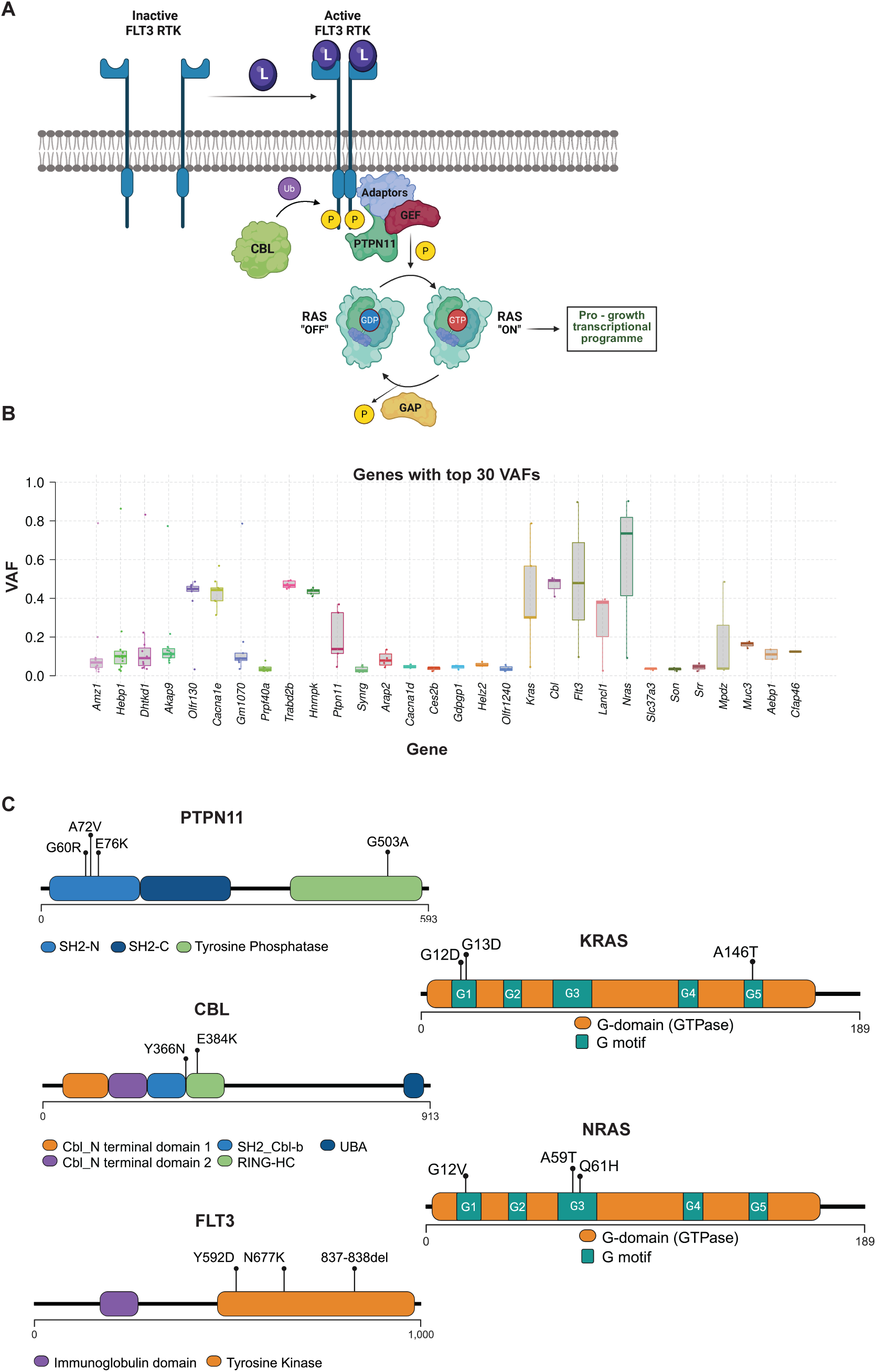
RTK-RAS pathway. **A.** RAS proteins are GTPases and cycle between "on"/"off" states. In the "off" state the protein is bound by GDP and in the "on" state by GTP that contains an extra phosphate group. Abbreviations: L, Ligand; Ub, Ubiquitin; P, Phosphate group. **B.** Variant allele frequency (VAF) for the top 30 most frequently mutated genes in *CEBPA* N321D tumours, arranged from most common to least common. VAF is calculated as the percentage of reads that do not correspond to the reference variant divided by total reads mapping to the region in a sample. **C.** Mutations in the RTK-RAS pathway are in functional protein domains that resemble those found in human disease. Lollipop plots of mutations found in the RTK-RAS pathway genes in the *CEBPA* N321D transformed tumours. The location of the mutations is shown alongside the functional domains of the proteins.

To find patterns amongst the observed mutations in the tumour samples, pathway enrichment analysis was performed. The most overrepresented pathway was the RTK-RAS pathway (present in 14/14 samples), which included mutations in *Ptpn11*, *Cbl*, *Flt3*, *Kras* and *Nras* (**Figure 3B**). Several samples contained multiple mutations in RTK-RAS pathway genes, although most had only a single mutation. The RTK-RAS pathway is central for cell growth, differentiation, and cell survival (**Supplementary Figure 5A**), with pro-oncogenic mutations found across a wide spectrum of human cancers ^16,17^. RTK-RAS mutations presented with high variant allele frequency (VAF), reaching >50% VAF in some samples hinting at the potential presence of homozygous clones. Moreover, RTK-RAS pathway mutations were amongst the highest calculated VAFs across all the mutations (**Supplementary Figure 5B**) and were commonly found within key functional domains of the respective protein products (**Supplementary Figure 5C**). Combined with their presence in every single tumour sequenced, these observations suggested that RTK-RAS pathway mutations represent pivotal late drivers of leukaemogenic progression in our model of CEBPA-mutated myeloid neoplasm.

Strikingly, most of the observed mutations in the RTK-RAS proteins recapitulated the exact mutations found in human patients (**Figure 3C**), including adult Acute Myeloid Leukaemia (AML) ^18,19^, paediatric AML and Acute Lymphoblastic Leukaemia (ALL) ^20^ and Juvenile Myelomonocytic Leukaemia (JMML) ^21,22^. Mutations in *RAS* (*KRAS*, *NRAS* and *HRAS*) are one of the most common mutations seen in human cancers, and are associated with poor prognosis and increased mortality ^16,17^. In particular, 98% of *RAS* mutations in human cancers are found at three amino acid residues: G12, G13 or Q61. In our dataset, we observed *KRAS* G12D, *KRAS* G13D, *NRAS* G12V and *NRAS* Q61H mutations, all of which have been reported as common *RAS* mutations across a variety of human cancers (Catalogue of Somatic Mutations in Cancer (COSMIC), reviewed in Muñoz-Maldonado *et al.*^23^). Exome sequence analysis therefore confirmed that the *CEBPA* N321D mutation is not sufficient to drive overt leukaemia, yet it imposes in this particular model a strong selection for *de novo* acquisition of RTK-RAS pathway mutations, that are an exact match to known oncogenic drivers across a large spectrum of human cancers.

### Leukaemic transformation is a late event during disease progression

So far, the *in vivo* experiments showed that (i) leukaemia develops with relatively long latency (median of 255 days after the *CEBPA* N321D cells were transplanted, without irradiation), (ii) all primary tumours contain mutations in the RTK-RAS pathway, and (iii) secondary transplants cause leukaemia with much reduced latency. This prompted us to investigate the temporal emergence of the transforming RTK-RAS pathway mutations. To differentiate between early-versus late-acquisition of RTK pathway mutations, *CEBPA* N321D cells were injected into murine recipients and three mice (m65, m68 and m71) were culled 3 months post-transplantation and their BM cells harvested. An aliquot of unfractionated BM cells was retained for subsequent molecular analysis, while the remainder cells were transplanted into secondary recipients (**Figure 4A**). Each of the three primary cases (‘donor cells’) was transplanted into at least three secondary recipient mice, and animals were monitored for leukaemia development. One secondary recipient from each group was culled 226 days after the primary transplantation (133 days post-secondary transplantation) as an intermediate sample (thereafter called ‘7 months intermediate’) whilst the remaining mice were left for tumour development.

**Figure 4.**
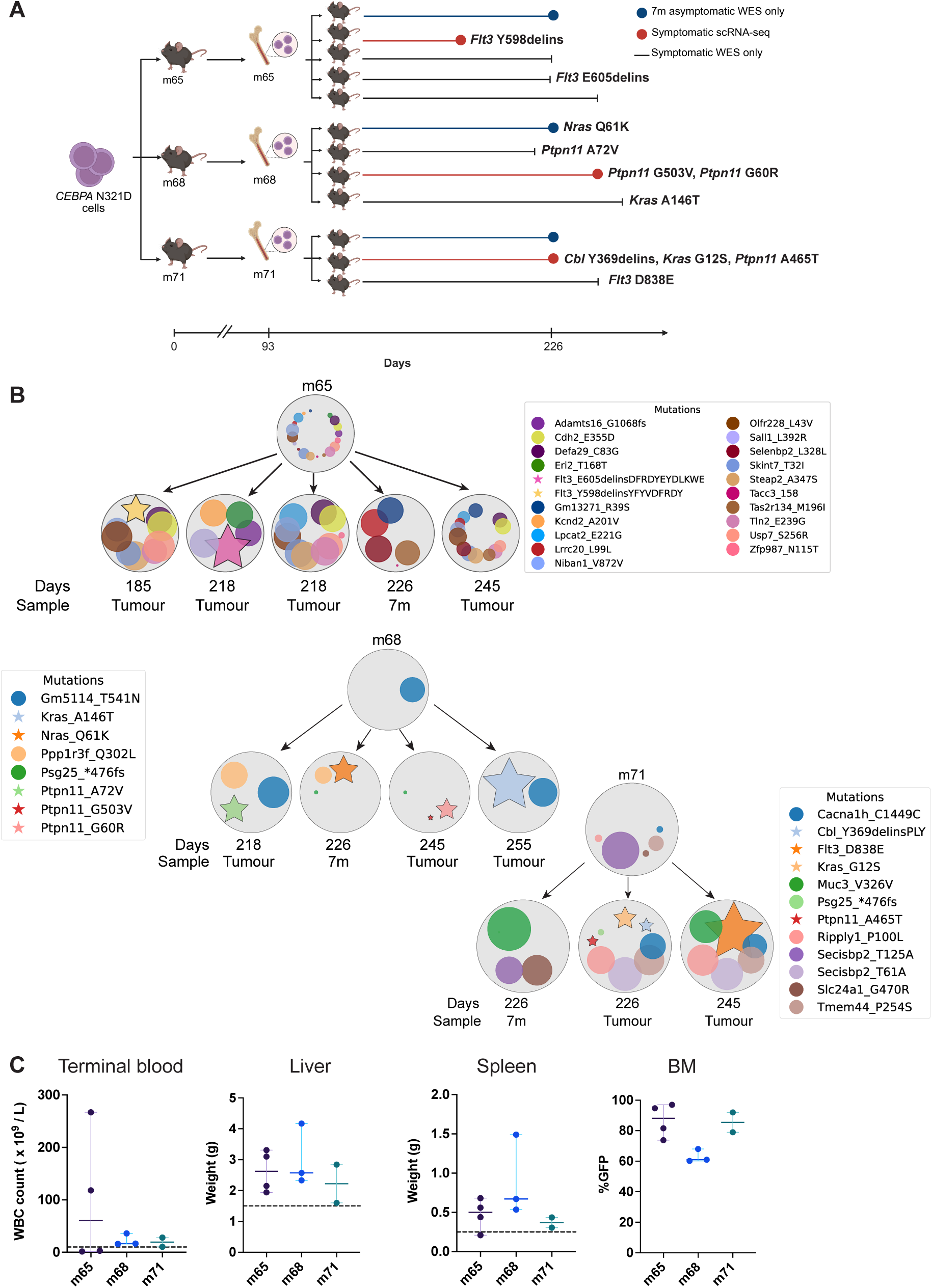
Leukaemia transformation is a late event during disease progression. **A.** Schematic of the experiment to determine the timeline of *CEBPA* N321D mediated tumorigenesis. *CEBPA* N321D cells were injected into three mice (m65, m71 and m68). After 93 days, these mice were culled, and their BM cells transplanted into secondary recipients. One mouse from each of the three groups was culled at 226 days post primary transplantation. The remaining animals were monitored for leukaemia development. The timeline at the bottom of the figure represents the survival duration of the secondary recipients, with RTK-RAS mutations identified in their BM cells annotated alongside the survival curves. Each sample is labelled with the number of days after the primary transplantation. BM cells from all samples were analysed by WES. Symptomatic samples submitted for 10x Genomics single-cell RNA-seq are highlighted in red. **B.** Schematic of mutations found in the BM cells of the 93-day donor and secondary recipients. Each passenger mutation is displayed as a circle and a RTK-RAS driver is shown as a star shape. The size of the shape is proportional to the VAF of that mutation. To simplify the schematic, we filtered out mutations that appeared at high VAFs in all samples. Additionally, for the m65 samples, only mutations that emerged from the 93-day donor sample onward are included. All shared mutations and their corresponding VAFs can be found in Supplementary Figure 6. **C.** Graphs show hepatomegaly and splenomegaly, increased WBCs in terminal bloods and high contribution of GFP-positive cells in symptomatic animals.

**Supplementary Figure 6.**
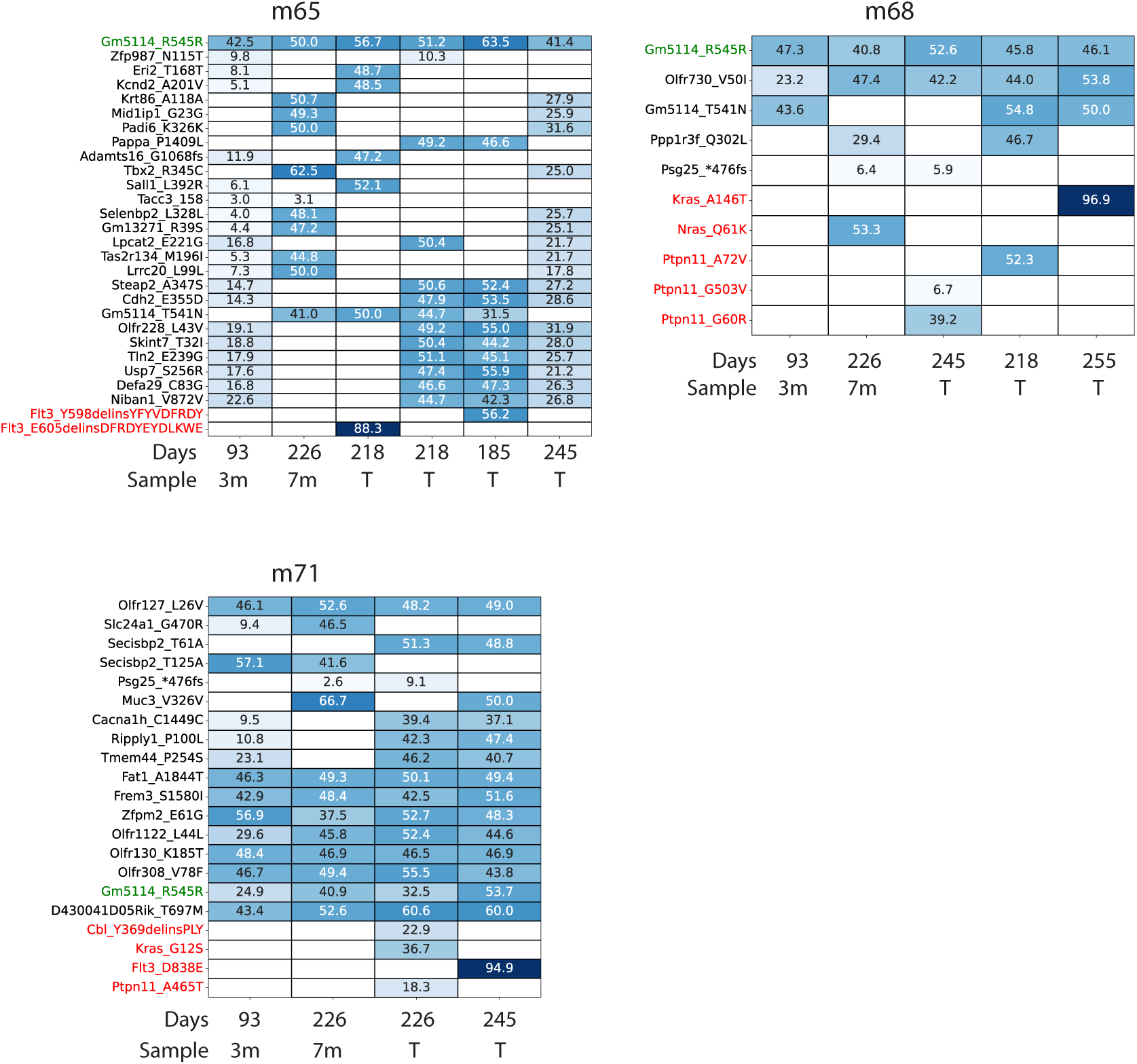
Full list of mutations in BM cells from mice injected with *CEBPA* N321D cells. Mice were culled 93 days post-injection ("donor cells") or, after transplantation of the donor BM cells into secondary recipients, either culled 4 months later (226 days intermediate) or monitored for the development of a leukaemia phenotype. Only mutations present in more than 2 samples per group are displayed. Each box shows the VAF of a mutation, with the blue colour intensity proportional to the VAF. Silent mutations are shown in green (no change in the coding amino acid). Mutations in RTK-RAS members are shown in red.

Interestingly, survival plots showed that tumours developed asynchronously amongst the recipients that received cells from the same donor at early time points (**Figure 4A**). Moreover, RAS-RTK mutations were only detected in the symptomatic mice, with none detected in the 3-month donor samples. One single exception was the 7-month mouse from the m68 donor group, which had no clinical signs of disease, but had a *Nras* Q61K mutation (VAF 53.3%) and a 33% GFP-positive population in the BM, compared to only 5% in samples lacking driver mutations. Therefore, we suggest that this sample captured an early transformation event; the acquisition of a RAS-RTK mutation that triggered malignant clone expansion but had not yet had time to generate overt leukaemia symptoms. Finally, all tumours displayed different driver mutations, suggesting independent mutation acquisition within the reconstituted marrow in the donor, yet interestingly the mutations were in the same five members of the RAS-RTK pathway as seen in previous experiments: *Flt3*, *Nras*, *Ptpn11*, *Kras* and *Cbl* (**Figure 4B**). Of note, three of the mice developed tumours with mutations with the exact amino acid changes found from the first transplantation experiment, namely *Ptpn11* A72V, *Ptpn11* G60R and *Kras* A146T, demonstrating a remarkable specificity towards the type of secondary mutation required to fully transform our model.

Disease was confirmed in all symptomatic secondary recipients by demonstrating typical signs of leukaemia, including leukocytosis, hepatomegaly, splenomegaly and a high proportion of mutant GFP-positive cells in the BM (**Figure 4C**). To further characterise the mutational dynamics of disease progression, we catalogued non-recurrent neutral passenger mutations that accumulated over time, as barcodes to infer the evolution of different clones within our samples. By tracking these mutations, we identified distinct clonal expansions in the individual mice from all three groups of secondary recipients (**Figure 4B, Supplementary Figure 6**), confirming that leukaemic transformation occurred independently in different clones. For example, two tumours that arose from m65 cells not only acquired distinct mutations in the *Flt3* gene but also exhibited the expansion of a different subset of passenger mutations. Similarly, a mutation in the *Gm5114* gene present in the 3-month donor cells for the m68 group, showed increased VAF in two tumours, yet was absent in the other two samples. Moreover, each secondary recipient acquired a unique mutation in the RTK-RAS pathway, underscoring independent clonal evolution. Finally, although two tumour samples from the m71 group shared four passenger mutations, the 7-month intermediate did not display these mutations and instead showed an expansion of two other passenger mutations from the 3-month sample. Collectively, these results are consistent with a model whereby multiple clones exist by the three-month time point, each with the potential to subsequently evolve into a tumorigenic clone.

### Overt leukaemia develops without a differentiation block where individual tumours produce either lymphoid or myeloid output

To further characterise the *CEBPA* N321D cells before and after transformation, we performed single-cell RNA sequencing (scRNA-Seq) on the same bone marrow aliquots used for exome-sequencing (**Figure 4A**). To focus the scRNA-Seq on the *CEBPA* N321D derived cells, we processed sorted GFP^+^ cells from all samples. We analysed (i) *CEBPA* N321D cells collected on the day of primary transplantation (“Preinjected” - day 0 Figure 4), (ii) cells from the m65, m68, and m71 mice at 93 days post-injection (“Asymptomatic”), and (iii) cells obtained from the secondary transplant recipients of the m65, m68, and m71 BM donor cells as the recipients were being sacrificed because of their full-blown leukaemia (“Symptomatic” - highlighted by red circles in Figure 4A). Additionally, we profiled GFP^-^ host progenitor cells, defined here as Lineage^-^ cKit^+^, from BM samples collected from the 93-day group (m65, m68, m71) to have a reference set of progenitors obtained from the same mice on the same day.

Evolution of cellular states during leukaemia development was assessed by projecting the new single cell transcriptomes onto a non-diseased mouse whole bone marrow (BM) atlas (Sturgess et al., in preparation), comprised of 196,986 cells isolated from the BM of non-diseased wild-type mice combining haematopoietic stem/progenitor enriched populations (lineage^-^ c-kit^+^ and lineage^-^ c-kit^+^ Sca1^+^) with total mononuclear bone marrow cells thus providing a comprehensive reference for non-diseased mouse haematopoiesis (**Figure 5A**). As expected, the 93-day host progenitor cells matched across a broad range of haematopoietic progenitor populations. These Lin^-^ cKit^+^ cells represent the endogenous recipient mouse haematopoietic system, thereby confirming the validity of our projection method (**Supplementary Figure 7**). The ‘pre-injected’ *CEBPA* N321D cells most closely resemble immature monocytes, particularly myeloid-monocyte progenitors (MMP3) and granulocyte-monocyte progenitor (GMP) populations (**Figure 5B**). Three months (93 days) after transplantation (**Figure 5Ci**), cells harvested from three individual recipient mice (m65, m68, and m71) appear highly similar to one another but map transcriptomically across a broader range of cell populations, compared to the pre-injected sample. In all three 93-day samples, there is a notable expansion of pDC-like cells and the emergence of lymphoid-like, immature B-cell populations.

**Figure 5.**
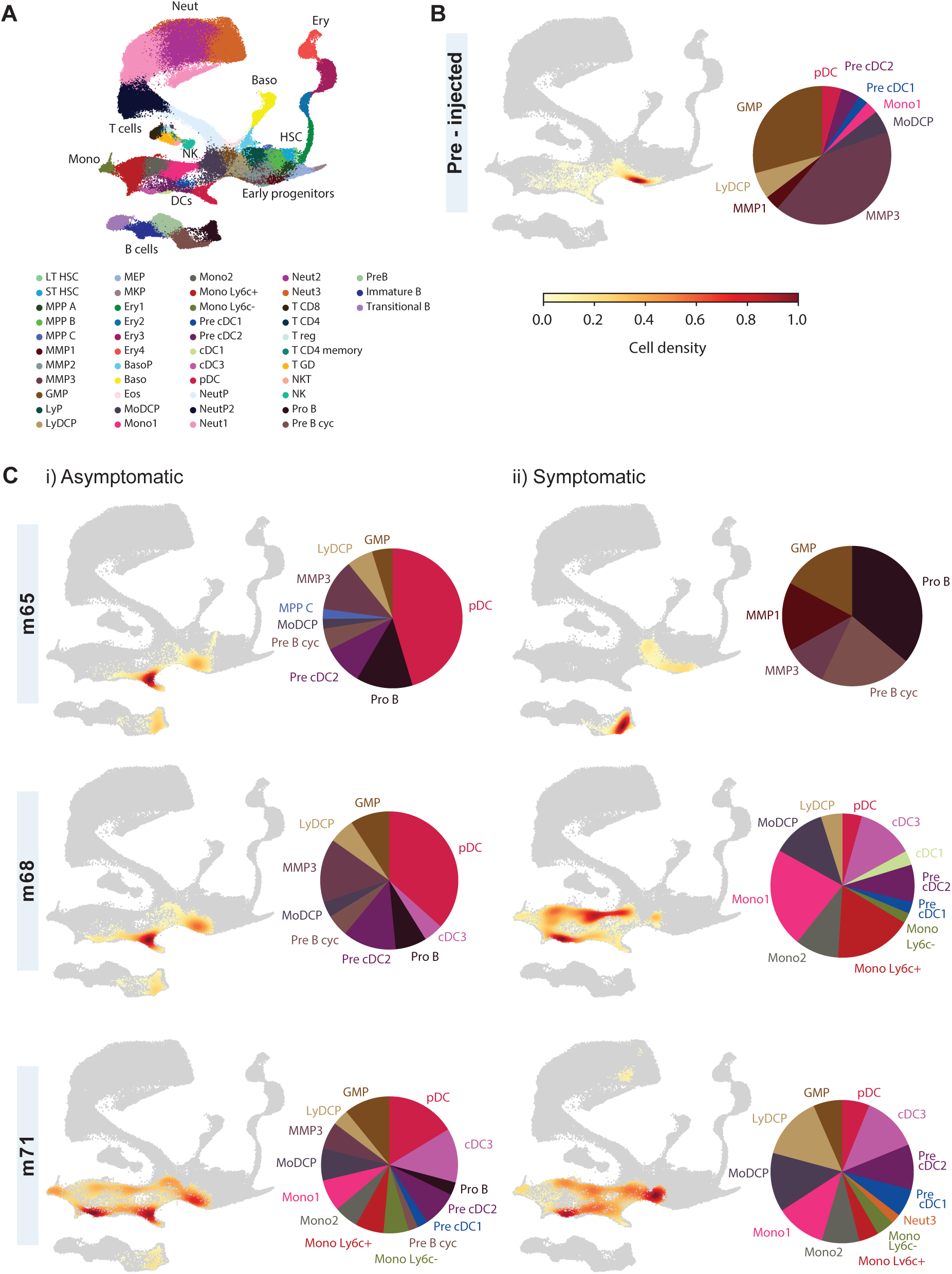
Leukaemogenesis in the *CEBPA* N321D model depends on the secondary mutation. **A.** Non-diseased mouse BM atlas used as a reference for projection and cell type annotation of our samples. Long term haematopoietic stem cell – LT HSC; short term haematopoietic stem cell – ST HSC; multipotent progenitor – MPP; myeloid-monocyte progenitor – MMP, granulocyte-monocyte progenitor – GMP; lymphoid progenitor – LyP; lymphoid dendritic cell progenitors - LyDCP; megakaryocyte-erythroid progenitors - MEP; megakaryocyte progenitor - MKP; erythrocyte - Ery; basophil progenitor – BasoP; basophil - Baso; eosinophil - Eos; monocyte-dendritic cell progenitors - MoDCP; monocyte - Mono; conventional dendritic cell – cDC; plasmacytoid dendritic cell – pDC; neutrophil - Neut; natural killer cell – NK. **B.** Results of cell type annotation of *CEBPA* N321D cells collected on the day of transplantation, ‘pre-injected’; **C.** Cell type annotation of i) Cells collected 93 days post primary transplantation from mice m65, m68 and m71 (asymptomatic); ii) symptomatic recipients of either the m65, m68 or m71 donor cells.

**Supplementary Figure 7.**
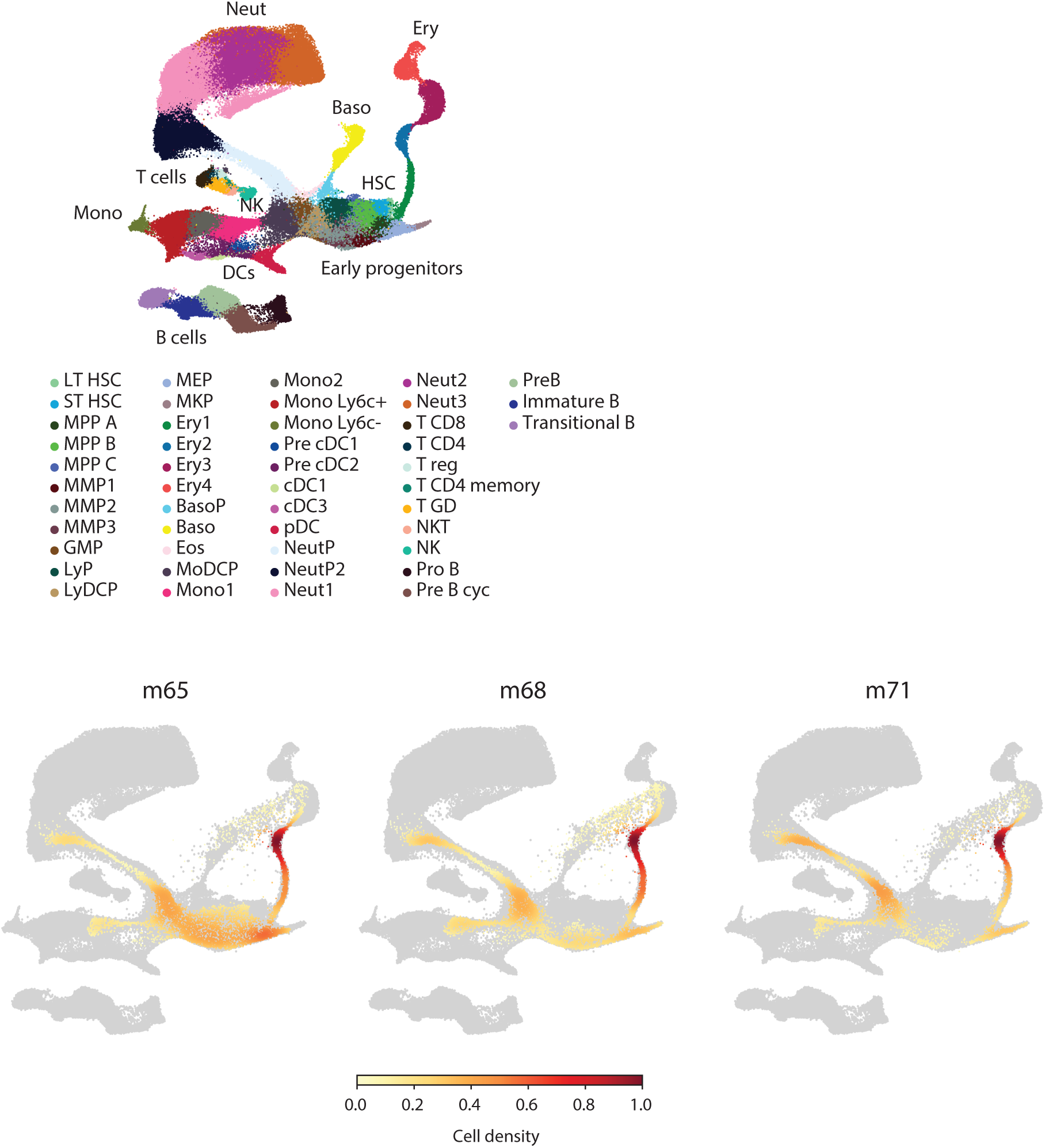
Results of cell type annotation of Lineage^-^ cKit^+^ cells sorted from the m65, m68 and m71 donors at the 93-day time point.

While all three asymptomatic samples (day 93) covered a similar range of cell states, the two symptomatic tumours derived from m68 and m71 donors mapped to immature myeloid progenitors and monocyte populations, whereas the tumour derived from the m65 donor mapped to immature lymphoid progenitors and B-lymphoid cells (**Figure 5Cii**). Cross referencing with the exome sequencing data outlined in the previous section showed that the m68 tumour contained two mutations in *Ptpn11*, the m71 tumour carried a different *Ptpn11* mutation along with additional mutations in *Cbl* and *Kras*, whereas the m65 tumour harboured a *Flt3* mutation. Single cell RNA-Seq thus showed that progression from preleukaemia to full-blown disease in this model is not associated with a strict differentiation block and accumulation of cells, since both stages are characterised by a differentiation axis going from immature to more mature cells. Moreover, overt leukaemia can adopt either a myeloid or lymphoid phenotype resulting in a somewhat perplexing situation whereby the model displays exquisite specificity for a particular pathway (RAS-RTK) for secondary mutations yet permits the development of leukaemias of different lineage phenotypes. Of note, both these features are reminiscent of Juvenile myelomonocytic leukaemia (JMML).

## Discussion

Understanding the mechanisms of oncogenic progression remains a major challenge in cancer biology, largely due to the difficulty of accessing the early stages of the disease in human patients. Given the parallels between relapse and the transition from premalignant to full blown cancer, a deeper understanding of leukaemogenic mechanisms therefore has major implications both for relapse prevention and treatment of established disease. In this study, we describe a novel tractable model of leukaemogenesis with characteristic acquisition of secondary RTK-RAS pathway mutations, that can be used to disentangle molecular and phenotypic cancer evolution *in vivo*.

Unlike conventional transplantation-based mouse models that use heterogeneous progenitor cells as their starting population, we took advantage of the Hoxb8-FL cell system that allows introduction of oncogenes into a well-defined progenitor population which maintains primary cell characteristics upon transplantation because of the switch-like nature of the Hoxb8-ER fusion transgene. We now demonstrate that retroviral introduction of a transgene carrying the C-terminal *CEBPA* N321D mutation removes the requirement for Hoxb8 for *in vitro* culture, leading to a pool of self-renewing progenitors with a pDC-like phenotype that can be propagated indefinitely with the supplementation of Flt3L. Transplantation of these differentiation-stalled cells *in vivo* showed them to be pre-malignant, requiring specific secondary mutations to develop full-blown disease with a latency of almost a year.

A particularly striking feature of the resulting disease was the exquisite specificity of secondary mutations, with all tumours analysed acquiring RTK-RAS pathway secondary mutations. It was furthermore intriguing to find that many of the mutations acquired in the endogenous mouse gene loci resulted in the exact same amino acid changes seen in human patients, including some of the most common RAS mutations found across human cancers. Within the haemato-oncology domain, mutations in *Kras*, *Nras*, *Ptpn11* and *Cbl* genes serve as one of a constellation of diagnostic markers for JMML, a rare but aggressive paediatric myeloid neoplasm, characterised by overactive RAS signalling ^28^. Other alterations in RTK-RAS pathway genes, such as *FLT3-*mutations, have also been described, albeit at a lower frequency ^29,30^. One of our tumours had acquired the most common JMML mutation *Ptpn11* E76K ^21^. Current JMML treatment strategies mainly involve allogeneic haematopoietic stem-cell transplantation (HSCT), but high relapse rates are observed, and the disease is often fatal.

Our *CEBPA* N321D preleukaemic cells demonstrated pDC-like characteristics. Human malignancies associated with pDC defects include the rare disease entities blastic plasmacytoid dendritic cell neoplasm (BPDCN) and AML with pDC expansion (pDC-AML), and neither have been particularly reported to associate with RAS specific mutational profiles. Of note, DC-like lineage capacity of JMML cells has previously been described ^31–33^. Another distinguishing feature of the leukaemias observed in this study is that they lacked the differentiation block commonly associated with myeloid malignancies such as AML. Strikingly JMML is characterised by a low percentage of blasts (<20%) ^34,35^. Moreover, two independent studies using flow cytometry or single-cell RNA sequencing have confirmed the presence of all major differentiating haematopoietic cell types in JMML human samples ^36,37^, demonstrating that JMML lacks the differentiation block that is commonly observed in acute leukaemias ^34^. Furthermore, our results showed that both myeloid and B-lymphoid output was produced during the preleukaemic phase, whereas full blown leukaemias produced output for just one of those downstream lineages. In rare case reports, individual JMML patients have been described in the clinic, with an abnormal B-cell population ^38^ and disease progression to a pre-B cell acute lymphoblastic leukaemia ^39^. Setting it apart from other mouse models, our RTK-RAS–driven model shows high specificity for secondary mutations, yet flexibility in terms of haematopoietic lineage output, all reminiscent of the disease seen in human JMML patients.

RAS proteins had been long thought of as undruggable although recent efforts have resulted in several KRAS inhibitors currently in Pre-clinical or Phase I clinical studies (reviewed in ^40,41^). Models of common RAS mutations that recapitulate human haematological cancers are therefore badly needed. The leukaemic cells generated in this study harbour patient relevant mutations and could serve as an attractive model of *RAS* mutated leukaemia. Tumour derived BM cells from our new model readily engraft into secondary recipients thus permitting *in vivo* testing of any new therapeutics, including the development of drug resistance. Unlike transgenic models, the mutations arise within the endogenous gene locus. Mutant RAS is therefore under the control of endogenous regulatory sequences, which is ideal to model resistance mechanisms driven by altering gene expression levels including epigenetic processes. Moreover, our model overcomes a key limitation of patient derived xenograft models (PDX), where patient derived cells are transplanted into immunocompromised animals that have been genetically engineered to permanently lack an immune system. This not only causes the loss of a functional tumour microenvironment, that is often essential in evaluating drug responsiveness but also makes it impractical to assess any combination therapies where new RAS inhibitors are combined with the rapidly growing array of immune-oncology therapeutics.

Although primary human tissues provide a valuable tool for studies into carcinogenesis, material is often limited. Moreover, *in vitro* cell culture from such samples can be extremely challenging, especially in malignancies that lack a full differentiation block, including JMML ^42^. Moreover, JMML, although potentially a devastating disease in children, is incredibly rare, thus models for its study are lacking. Here we report a new model of leukaemogenesis that provides access to the full timeline of tumour development, thus enabling direct comparisons across all stages of malignant transformation, to not only develop hypotheses for subsequent mechanistic investigation, but also to test the efficacy of new therapeutic approaches. With regards to the latter, our model stands out by being fully immunocompetent and developing patient relevant RAS-RTK mutations within the endogenous gene loci, thus providing a robust preclinical platform to accelerate the development of precision therapies, bridging critical gaps between mechanistic discovery and clinical translation.

## Methods

### Cell lines and culture conditions

Hoxb8-FL cell line was kindly provided by the Hans Häcker laboratory. Cells were grown at 37°C and 5% CO_2_ in RPMI 1640 medium (Sigma R8758) supplemented with 10% Foetal Bovine Serum (FBS) (Gibco 10270-106), 5% Flt3L conditioned medium, 0.1% 2-Mercaptoethanol (Invitrogen 31350-010), 1% Penicillin/ Streptomycin (Sigma, P0781), 1% Glutamine (Sigma, G7513) and 1μM β -oestradiol (Sigma E2758). This media is termed as self-renewal media (Flt3L+Oe+) throughout the text. Cells were kept at densities between 0.7 - 12 x 10^5^ cells/mL.

Flt3L conditioned medium was prepared using the B16-FLT3 cell line that constitutively expresses Flt3L and was kindly provided by Hans Häcker laboratory. The B16 cells were seeded in 10 cm dishes and grown at 37°C and 5% CO_2_ in RPMI 1640 medium (Sigma R8758) supplemented with 10% FBS (Gibco 10270-106), 0.1% 2-Mercaptoethanol, 1% Penicillin/ Streptomycin (Sigma, P0781), 1% Glutamine (Sigma, G7513). Cells were cultured for 2-3 days until confluent, the supernatant harvested, and media replaced daily over a 3-day period. The supernatant was then filtered, aliquoted and stored at −80°C.

293T cells were cultured in Dulbecco’s Modified Eagle Medium (DMEM) supplemented with 10% FBS, 1% Penicillin/ Streptomycin (Sigma, P0781) and 1% Glutamine (Sigma, G7513). Cells were cultured in 100 mm/150 mm dishes at 37°C and 5% CO_2_. Cells were passaged every 2-3 days with Trypsin-EDTA (Sigma, T3924) and kept at 30-90% confluency.

Differentiation media refers to the Flt3L media that induced differentiation of Hoxb8-FL cells and consisted of the same components that the Hoxb8-FL media except without the addition of β-oestradiol.

### Cloning of the retroviral *CEBPA* N312D, *CEBPA* WT and Empty Vector (EV) plasmids

The pMSCV-IRES-eGFP (EV), pMSCV-WT (human) CEBPA – IRES – eGFP (CEBPA WT) and pMSCV – N321D (human) CEBPA – IRES – eGFP (*CEBPA* N321D) constructs were generated as follows.

The human CEBPA N321D construct was assembled from synthetic nucleotides (GeneArt Life Technologies) and inserted into a kanamycin-resistant vector to generate pMK-T CEBPA N321D. The sequence for the inserted human N321D version of the *CEBPA* gene is the following: ATGGAGTCGGCCGACTTCTACGAGGCGGAGCCGCGGCCCCCGATGAGCAGCCACCTGCAGAGCCCCCCGCACGCGCCCAGCAGCGCCGCCTTCGGCTTTCCCCGGGGCGCGGGCCCCGCGCAGCCTCCCGCCCCACCTGCCGCCCCGGAGCCGCTGGGCGGCATCTGCGAGCACGAGACGTCCATCGACATCAGCGCCTACATCGACCCGGCCGCCTTCAACGACGAGTTCCTGGCCGACCTGTTCCAGCACAGCCGGCAGCAGGAGAAGGCCAAGGCGGCCGTGGGCCCCACGGGCGGCGGCGGCGGCGGCGACTTTGACTACCCGGGCGCGCCCGCGGGCCCCGGCGGCGCCGTCATGCCCGGGGGAGCGCACGGGCCCCCGCCCGGCTACGGCTGCGCGGCCGCCGGCTACCTGGACGGCAGGCTGGAGCCCCTGTACGAGCGCGTCGGGGCGCCGGCGCTGCGGCCGCTGGTGATCAAGCAGGAGCCCCGCGAGGAGGATGAAGCCAAGCAGCTGGCGCTGGCCGGCCTCTTCCCTTACCAGCCGCCGCCGCCGCCGCCGCCCTCGCACCCGCACCCGCACCCGCCGCCCGCGCACCTGGCCGCCCCGCACCTGCAGTTCCAGATCGCGCACTGCGGCCAGACCACCATGCACCTGCAGCCCGGTCACCCCACGCCGCCGCCCACGCCCGTGCCCAGCCCGCACCCCGCGCCCGCGCTCGGTGCCGCCGGCCTGCCGGGCCCTGGCAGCGCGCTC AAGGGGCTGGGCGCCGCGCACCCCGACCTCCGCGCGAGTGGCGGCAGCGGCGCGGGCAAGGCCAAGAAGTCGGTGGACAAGAACAGCAACGAGTACCGGGTGCGGCG CGAGCGCAACAACATCGCGGTGCGCAAGAGCCGCGACAAGGCCAAGCAGCGCAACGTGGAGACGCAGCAGAAGGTGCTGGAGCTGACCAGTGACGATGACCGCCTGCGCAAGCGGGTGGAACAGCTGAGCCGCGAACTGGACACGCTGCGGGGCATCTTCCGCCAGCTGCCAGAGAGCTCCTTGGTCAAGGCCATGGGCAACTGCGCGTGA

To generate the WT version of CEBPA, the pMK-T CEBPA N321D construct was digested with BlpI and BstAPI restriction enzymes (New England Biolabs) to excise a portion of CEBPA gene containing the mutated nucleotide. The resulting linearised product was run on an agarose gel and extracted with the QIAquick Gel Extraction Kit (Qiagen). Complementary primers containing the WT version of human CEBPA gene were annealed together and used to replace the region in the digested vector with the WT version of the gene.

To generate the retroviral constructs used in this study, the pMSCV-MLL-ENL-IRES-eGFP vector (obtained from S Basilico^12^ was digested with Sal I and Bgl II restriction enzymes (New England Biolabs) to release the MLL-ENL fragment. The rest of the fragment was either re-ligated to generate the control pMSCV-IRES-eGFP (EV) plasmid or used to insert the CEBPA gene in the following manner. The CEBPA WT or CEBPA N321D cDNAs were first amplified by polymerase chain reaction (PCR) using KAPA HiFi HotStart ReadyMix (Kapa Biosystems, now Roche). PCR primers were designed to attach an in-frame FLAG tag and Sal I / Bgl II restriction sites to CEBPA inserts. The resulting CEBPA WT/N321D-Flag inserts were ligated into digested pMSCV-IRES-eGFP vector by Gibson assembly and transformed into electrocompetent DH-10β cells. Ampicillin-resistant colonies were selected by growth on agar plates with ampicillin. Plasmid DNA was isolated by an alkaline lysis method using buffers S1 (containing RNaseA), S2 and S3 from the NucleoBond®Xtra Maxi Kit (Macherey-Nagel). The resulting plasmids verified by Sanger Sequencing.

### Retroviral production and transduction of Hoxb8-FL cells with CEBPA N321D, WT or EV

5 x 10^6^ 293T cells were seeded into 10 cm culture dishes approximately 16 hours before transfection, aiming for 80% cell confluency at the time of addition of the viral supernatant. 5 μg of DNA (MSCV-EV-GFP/MSCV-CEBPA-WT-GFP/MSCV-CEBPA-N321D-GFP) was mixed with 5 μg of Psi packaging vector, topped up to 10 μL with H_2_O and added to 500 μL of plain DMEM (DNA/DMEM mix). In another tube, 500 μL of plain DMEM was mixed with 45 μL of TransIT-LT1 Transfection Reagent (Mirus MIR 2305, TransIT/DMEM mix). The DNA/DMEM mixture was slowly added to the TransIT/DMEM mix and incubated at room temperature for 30 mins. The resulting solution was slowly distributed in a drop wise fashion around the dish of 293T cells and cells incubated overnight at 37°C and 5% CO_2_. 24 hours post transfection, the 293T media in the dishes was replaced with 6 mL of self-renewal Hoxb8-FL cell media and cells were incubated at 37°C and 5% CO_2_. After a further 24 hours, the cell supernatant was collected with a 5 mL syringe, taking extra care to not disturb the attached cells at the bottom of the dish. The collected viral supernatant was passed through a 0.45 μm filter to remove cell debris.

For transduction, 1.5 x 10^6^ of Hoxb8-FL cells were placed in 2 mL of appropriate media in a well of a 6-well cell culture plate. 24 μL of 1 mg/mL polybrene (Sigma TR-1003-G) was added to the well to a final polybrene concentration of 8 μg/mL. 1 mL of viral supernatant (carrying either MSCV-EV-GFP, MSCV-CEBPA-WT-GFP, MSCV-CEBPA-N321D-GFP) was added to the well and gently mixed by pipetting up and down and swirling of the plate. Cells were centrifuged at 780 x g, at 32°C for 90 mins (Allegra 6R Rotor GH-3.8) with a "No break" setting. Following the spin, cells were incubated for 90 mins at 32°C and 5% CO_2_. Subsequently, 1.5 mL of media was carefully removed from the side of the well (without disturbing the cells) and 2.5 mL of fresh media was added. Cells were incubated overnight at 37°C and 5% CO_2_.

### Differentiation conditions to generate the *CEBPA* N321D cells

Hoxb8-FL cells were transduced with either empty vector (EV), *CEBPA* WT or *CEBPA* N321D viral vectors and cultured for two days in self-renewing media (media used to culture the parental Hoxb8-FL cell line). Two days post transduction, cells were washed two times with phosphate buffered saline (PBS) (Sigma D8537) and resuspended in differentiation media (Flt3L) (RPMI 1640 medium (Sigma R8758) supplemented with 10% Foetal Bovine Serum (FBS) (Gibco 10270-106), 5% Flt3L conditioned medium, 0.1% 2-Mercaptoethanol (Invitrogen 31350-010), 1% Penicillin/ Streptomycin (Sigma, P0781) and 1% Glutamine (Sigma, G7513)). Cells were counted using the CASY® Cell Counter, counting cells with diameter range of 7.85 μm - 14.95 μm. Cells were seeded at a concentration of 1 x 10^5^ cells/mL in 6-well plates (2 mL total volume) and in T75 flasks (12 mL total volume) and cultured at 37°C and 5% CO_2_. Medium was changed every 2-3 days, and the cells were kept at a concentration of 0.7 - 16 x 10^5^ cells/mL. Differentiated cells that were adhered to the wells/flasks were dissociated enzymatically with Trypsin-EDTA (Sigma T3924) prior to processing. However, CEBPA N321D cells past Day 35 were not trypsinised as cells were no longer adherent. Presentation of haematopoietic markers was analysed by flow cytometry on the day cells were transferred into differentiation media (day 0) and at 5, 9, 14, 21 and 28 days after. The cells in plates were used for flow cytometry analysis at days 0, 5 and 9, the cells in flasks were used for all the subsequent flow cytometry characterisations and cell expansion. In the case of the CEBPA WT and EV cells, cells were monitored for 12 days until all cells differentiated and the rate of cell death precluded the continuation of the experiment. For CEBPA N321D cells, the cells were cultured for 35 days and cells frozen in 7 x 10^6^ cells/mL aliquots in 1 mL of 10% DMSO/90% FBS solution.

### Flow cytometry and sorting

Cells were centrifuged at 300 x g for 5 mins, washed two times with 2% FBS/PBS solution (Gibco 10270-106, Sigma D8537) and resuspended in 100 μl of 2% FBS/PBS. Cells were stained with an antibody mixture (see table below) and incubated in the dark at 4°C for 30 mins. Single stained controls were prepared using UltraComp eBeadsTM Compensation Beads (Thermo Fisher 01-2222-41). After incubation, cells were washed twice in 2% FBS/PBS, resuspended in 2% FBS/PBS and 1:175 dilution of 7-Aminoactinomycin D (7AAD) viability dye (BD 559925) was added to exclude dead cells. The LSR Fortessa (BD Biosciences) was used to analyse the cells. Antibodies used are listed below.

| Antibody | Clone | Supplier | Catalogue | Dilution |
| --- | --- | --- | --- | --- |
| c-kit (APC/Cy7) | 2B8 | BioLegend | 105826 | 1:100 |
| Ly-6G (BV510) | 1A8 | BioLegend | 127633 | 1:100 |
| B220 (AF700) | RA3-6B2 | BioLegend | 103232 | 1:100 |
| Cd11b (BV711) | M1/70 | BioLegend | 101242 | 1:800 |
| Cd11c (PE/Cy7) | N418 | BioLegend | 117318 | 1:100 |
| SIRPα (PE) | P84 | BD Biosciences | 560107 | 1:100 |
| XCR1 (APC) | ZET | BioLegend | 148206 | 1:100 |
| BST2 (BV421) | 927 | BD Horizon | 566431 | 1:100 |
| F4/80 (BUV395) | T45-2342 | BD Horizon | 565614 | 1:100 |
| MHC Class II (PerCP/Cy5.5) | M5/114.15.2 | BioLegend | 107626 | 1:400 |
| CD45.1 (PE/Cy7) | A20 | BioLegend | 110730 | 1:100 |
| CD45.2 (APC/Cy7) | 104 | BioLegend | 109824 | 1:100 |
| CD45 (BV711) | 30-F11 | BioLegend | 103147 | 1:100 |

In case of cell sorting, cells were sorted by the CIMR Flow Core Facility using the Influx Cell Sorter (BD Biosciences). Flow cytometry data was processed with the FlowJo software.

### Microscopy

For morphological examination of EV and *CEBPA* N321D cells, cytospin preparations were carried out by loading 50–100 µL of cells into cytofunnels and centrifuging at 700 rpm for 2 minutes on a Shandon Cytospin 3. The resulting cytoslides were stained using a Rapid Romanowsky Stain Pack (TCS Biosciences, HS705) according to the manufacturer’s protocol. Microscopic images were acquired on a Zeiss Axio Imager.Z2 using Zen Pro software.

Terminal blood samples were used to make blood smear slides, which were subsequently stained for haematoxylin followed by staining with eosin (H&E). Spleen, liver and bone marrow from symptomatic animals were preserved in formalin until submitted for processing and H&E staining to Cambridge Stem Cell Institute Histology facility. Images were taken on Zeiss Apotome 2.

### Submission of mouse bone marrow cells for 10X Genomics Single Cell RNA-Sequencing

30,000-60,000 mouse bone marrow cells were FACS sorted for GFP+ 7-AAD- into 2% PBS/FBS solution. Cells were washed with 2% PBS/FBS solution and resuspended in 0.04% Bovine Serum Albumin (BSA, AMBION) solution at concentration of 371 cells/μL. Cell were submitted to the Cancer Research UK Cambridge Institute Single-Cell Facility for 10x Genomics single cell RNA-Sequencing using the Chromium Single Cell 3’ Reagents (v3.1 Chemistry), libraries were prepared following manufacturers protocols and sequenced on an Illumina NovaSeq 6000.

### DNA extraction and preparation of material for whole exome sequencing

DNA from bone marrow cells and the *CEBPA* N321D cells was extracted using the Qiagen DNeasy Blood and Tissue Kit according to the manufacturer’s instructions for ‘Purification of Total DNA from Animal Blood or Cells (Spin-Column Protocol)’. Around 2 - 5 x 10^6^ cells were used for the extraction. The optional step of addition of 4 μl RNase A (100 mg/ml) was followed. DNA concentration was measured using the DeNovix Spectrophotometer. RNA contamination was checked by agarose gel electrophoresis. Exome capture and sequencing was performed by Dr. David Adams’ laboratory at the Wellcome Trust Sanger Institute or by Novogene (UK). Whole exome sequencing was performed with the Illumina Novaseq6000 platform (Wellcome Trust Sanger Institute) or with the Novaseq X Plus Series (Novogene).

### Sample processing for low-cell number RNA sequencing

Samples were processed using a modified version of the Smart-Seq2 protocol ^43,44^ described in Kucinski *et al* ^45^ with slight modifications. Briefly, cells were sorted into 96-well plates. For each well, 375 cells were sorted into 11.5 μl lysis buffer containing 1.15 μl SUPERase-In RNase Inhibitor (20 U/μl Thermo Fisher Scientific AM2696), 0.115 μl 100 mM DTT, 0.23 μl 10% Triton X-100 solution (Sigma 93443) and 10.005 μl Nuclease-free H_2_O (Sigma W4502). Cells were vortexed and spun in a pre-cooled centrifuge for 2 minutes at 2000 rpm and stored at −80 °C.

Plates were thawed on ice and 5 μl of annealing mixture added (0.5 μl External RNA Controls Consortium (ERCC) RNA Spike-In (1:300,000 dilution, Thermo Fisher 4456740), 0.1 μl Oligo-dT 100 μM primer and 4.4 μl Nuclease-free H_2_O (Sigma W4502)). Plates were centrifuged at 700 x g for 1 minute, incubated at 72°C for 3 minutes in a thermocycler and immediately cooled down on ice. 3.3 μl (or 1/5 of the total volume) of annealed material was transferred to a new plate and used for further processing. 6.7 μl of reverse transcription solution was added, containing 0.1 μl Maxima H Minus (200 U/μl, Thermo Fisher EP0753), 0.25 μl SUPERase-In RNase Inhibitor (20 U/μl Thermo Fisher Scientific AM2696), 1 μl dNTPs (10mM stock), 2 μl 5x Maxima RT buffer (Thermo Fisher EP0753), 0.2 μl TSO (100 μM, Qiagen), 1.875 μl PEG800 (40% v/v) (Sigma-Aldrich) and 1.275 μl Nuclease-free H_2_O (Sigma W4502). Samples were incubated at 42°C for 90 minutes and followed by an incubation at 70°C for 15 minutes. The cDNA was amplified by PCR. 40 μl of PCR mix was added consisting of 1 μl Terra PCR Direct Polymerase (1.25 U/μl, Takara 639271), 25 μl 2x Terra PCR Direct Buffer, 1 μl ISPCR primer (10 μM) and 13 μl Nuclease-free H_2_O (Sigma W4502). The PCR conditions were the following: 98°C for 3 mins (1 cycle); 98°C for 15 sec, 65°C for 30 sec, 68°C for 4 min (13 cycles); 72°C for 10 min (1 cycle). The PCR product was purified using AMPure XP beads (Beckman A63882).

Sequencing libraries were prepared as follows. 1.63 μl of Tagment DNA buffer, 0.60 μl of Tagmentase (Nextera XT DNA Library Preparation Kit, Illumina Fc-131-1096) and 1.03 μl cDNA (previously diluted to final concentration of 100-150 pg) were aliquoted into a new 96-well PCR plate and incubated a 55°C for 10 mins in a thermocycler. Immediately after incubation 0.82 μl of 0.2% SDS was added to neutralise the reaction. 1.23 μl of N Index Primer, 1.23 μl of S Index Primer 2 (Illumina Nextera XT Index Kit v2 Sets A (FC-131-2001) and D (FC-131-2004)) and 4.97 μl of PCR Master Mix (2.30 μl Phusion HF Buffer (Thermo Fisher), 0.10 μl dNTP (25mM stock), 0.07 μl, Phusion Polymerase, 2.50 μl (2 U/μl Thermo Fisher F530L), Nuclease Free H_2_O (Sigma W4502)) were added and PCR run with the following protocol: 72°C for 3 mins (1 cycle); 98°C for 15 sec, 65°C for 30 sec, 68°C for 4 min (13 cycles); 72°C for 10 min (1 cycle).

Library pool was cleaned with AMPure XP beads (Beckman A63882) and eluted in 35 μl EB (10mM Tris-Cl, pH 8.5; Qiagen 19086). Library size distribution was checked with High Sensitivity D5000 ScreenTape (Agilent 5067-5592). Libraries were quantified with KAPA qPCR quantification kit (Roche 07960140001/KK4824) using the MX3000P qPCR System (Stratagene, now Agilent). The pooled library was sequenced using the Illumina HiSeq 4000.

### Animals

Animal work was conducted using C57BL/6J mice. Animals were housed in pathogen-free animal facility in individually ventilated cages under a 12:12h light dark cycle, with food and water available ad libitum. Experiments were conducted under a UK Office project license, as per UK Home Office regulations. The animal work was performed in accordance with the Animals (Scientific Procedures) Act 1986, UK and was reviewed by the University of Cambridge Animal Welfare Ethical Review Body.

### *In vivo* transplantations

For all transplantation studies, 5 x 10^5^ of CD45.1+ cells were mixed with 1 x 10^5^ of CD45.1/2 helper cells and transplanted into lethally irradiated CD45.2 mice via tail vein injection. After transplantation, peripheral blood was withdrawn and chimerism was checked by measuring the levels of CD45.1, CD45.2 and green fluorescent protein (GFP) by flow cytometry analysis. Disease onset was also monitored by measuring white blood cells using a Blood Cell Counter Machine (scil Vet abc PlusTM). The specifics for each *in vivo* experiment are outlined below.

#### (a) Injection of Hoxb8-EV and CEBPA N321D cells into mouse recipients

For primary tumour induction experiments, either EV or *CEBPA* N321D cells were used. Prior to transplantation, to ensure a homogeneous population of cells, EV cells were FACS sorted for GFP+ cells and grown in self-renewal (Flt3L+Oe+) medium whereas *CEBPA* N321D cells were confirmed to be 100 % GFP+ via flow cytometry and grown in Flt3L+ medium. 20 four-week-old female mice were injected, with 17 mice receiving the mutant *CEBPA* N321D cells and 3 mice transplanted with EV cells.

#### (b) Transplantation of primary tumours into secondary mouse recipients

Two primary tumours which developed in mice from the experiment described in (a) were chosen. Non-sorted, bone marrow cells were injected into 5 mice per tumour. Both male and female recipient mice were used. Mice were between 8 - 14 weeks old.

#### (c) Transplantation of cells for the Mutational Time course experiment (as shown in Figure 4) CEBPA

N321D cells were injected into 8 nine-week-old female mouse recipients. Simultaneously, an aliquot of 60,000 CEBPA N321D cells was submitted for 10x Genomics single cell RNA-sequencing, ‘pre-injection’ timepoint. Instead of waiting for tumour development, 3 mice (cases) were culled 3 months after cells were injected and 20,000 cells from each case were submitted for 10x Genomics single cell RNA sequencing ‘3-months’ timepoint. The remaining cells from each case were re-transplanted into new lethally irradiated recipients, with at least 3 mice injected with 500,000 total bone marrow cells per case. No control cells (EV), were injected in this round. From each of the three cases, mice were culled 4 months post-transplantation, ‘7-months’ timepoint. Mice displaying signs of leukaemia development, including elevated white blood counts, were culled the bone marrow and spleen isolated, checked for percentage of GFP expression via flow cytometry and material frozen. Three samples of bone marrow cells from diseased mice were submitted for 10x Genomics single cell RNA-sequencing, where each of the three samples were recipients of one of the 3 cases from the ‘3-months’ timepoint.

### Bioinformatics

#### Generation of a custom reference file

Unless otherwise stated, the following reference mouse genome file was used for the alignment of sequencing data. FASTA file containing the sequence information and gene annotation GTF file for Mus musculus GRCm38 (mm10) was downloaded from ENSEMBL version 93. Genome sequence and annotation were then augmented with the sequence details of the 92 ERCC RNA spike-in controls, pMSCV plasmid backbone, GFP and the portion of human *CEBPA* sequence with the N321D mutation used in the generation of the MSCV-CEBPA-N321D-GFP plasmid.

### Analysis of 10x Genomics single-cell RNA-sequencing data

Read processing and alignment to the modified mm10 reference genome were performed using the 10x Genomics Cell Ranger Software Suite (version 7.0.1). Downstream analysis was performed with Scanpy framework^46^and functions from CellProject (https://github.com/Iwo-K/cellproject) with the singularity container from Kucinski *et al*.^47^. For quality control, cells with less than 500 genes or less than 5,000 counts were excluded. Genes not detected (zero counts across all cells) were removed. Potential doublets were detected using Scrublet^48^ (expected doublet rate = 10%) and removed from the dataset. Data were normalised to a total of 10,000 counts per cell, followed by log-transformation (log1p).

To annotate cell types in the query dataset, a non-diseased bone marrow reference atlas was used in combination with functions from the CellProject package (as described in Kucinski *et al*. ^47^). To reduce computational load, the reference was subsampled to a maximum of 1,000 cells per cell type. Log-normalised counts from the query and reference datasets were concatenated, and batch effects were removed using Seurat’s CCA method^49^ using only the highly variable genes from the reference. The reference data were scaled, and the first 50 principal components were computed. The corrected query values were then fit into the computed PCA space, and 15 nearest neighbours were identified between query and reference cells. These neighbours were used both to transfer cell labels (by majority vote) and to predict query coordinates in the reference PCA space, which were subsequently used to embed the query data into the reference UMAP space^50^. For visualization, a small subset of cells (760 out of 104,620) with a mitochondrial content greater than 7.5% was excluded. Additionally, only cells assigned to groups representing more than 2% of the total cells in each sample were retained. Finally, cell density in the reference UMAP space was calculated for each sample using Scanpy’s ^46^ embedding_density function (scanpy.tl.embedding_density) and visualised.

### Whole exome sequencing analysis

Forward and reverse reads were aligned with Burrow – Wheeler Aligner (BWA) mem algorithm (version 0.7.17) ^51^ using the custom reference mouse genome. The index for the reference file used for the alignment was created by BWA index ^51^. For quality processing of aligned reads, tools available from the GATK (version 4.3.0) suite ^52^ were used. Aligned SAM files were sorted by coordinate with SortSam (Picard) tool. PCR duplicates were marked by MarkDuplicates (Picard). Next, the resulting files were processed to improve the accuracy of the sequencing assigned base quality scores. First, the recalibration table was calculated with the BaseRecalibrator tool (GATK) by supplying the Mouse Genomes Project SNP and indel database (Release Version 5) as reference data for sites of known variation. Secondly, the resulting table was used to recalibrate the base quality scores of the input reads with the ApplyBQSR tool (GATK). The recalibrated files were used to detect short somatic variants with Mutect2 (GATK). Detected mutations include single nucleotide changes, small insertions and deletions. The analysis was run for each tumour sample against a matched normal, where the normal sample was the preinjected *CEBPA* N321D cell line. Low quality variants were flagged using FilterMutectCalls (GATK) with default parameters. Variants that passed the filtering were filtered into a new file with BCFtools (1.9) view command^53^. Variant annotation was done using the ANNOVAR package^54^. Mouse gene definition database was created for the mm10 mouse genome following ANNOVAR’s documentation by first downloading the UCSC mm10 annotation database via annotate_variation.pl -downdb and then generating a corresponding FASTA file with annotate_variation.pl –buildver and re-trieve_seq_from_fasta.pl generating a FASTA file with 47,382 mouse mm10 genomic regions. To generate a gene-based annotation the VCF files containing unannotated variant regions (from the ‘Variant calling’ step) were converted to annovar objects with convert2annovar.pl with -allsample –include –comment arguments. The resulting files were used to annotate the variants to genes via annotate_variation.pl –geneanno and default parameters. The resulting annovar objects was converted to maf files with the maftools package (version 2.12.00) ^55^ and visualisations created with maftools in R and custom code in Python.

### Analysis of low-cell number Smart-Seq2 RNA sequencing data

Sequencing reads were aligned to a modified mm10 reference genome (described above) using STAR aligner, and gene counts were quantified with featureCounts. Further analysis was performed using the in-house smqpp package (https://github.com/SharonWang/smqpp). Cells were filtered out if they met any of the following criteria: fewer than 100,000 reads; fewer than 4,000 highly abundant genes (detected above 10 counts per million); fraction of gene-derived counts relative to total counts below 15%; more than 12% ERCC-derived counts; or more than 5% mitochondrial-derived counts. Genes detected in fewer than 1 cell were filtered out and remaining counts were normalised using the smqpp.normalise_data function based on DESeq2^56^ size factor normalisation. The data were log-transformed, and 1,131 highly variable genes were identified. The data were then scaled, and PCA was performed on the expression values of the highly variable genes, retaining 50 principal components. A k-nearest neighbour graph was computed using 15 neighbours and the first 40 principal components, and UMAP embedding was generated.

## Figure generation

The authors would like to acknowledge that Figures 1A, 2A, 3A, 4A and Supplementary Figures 5A and 5B were created with BioRender.com.

## Data and code availability

All sequencing files are available in the GEO repository under the SuperSeries: GSE300817. This study did not generate any original code. Example notebooks of data analysis are available here: https://github.com/MariaZaro/StepwiseMyeloidModel/

## Author contributions

M.Z. designed and performed experiments, carried out bioinformatic analysis and wrote the manuscript; M.Q. designed and performed experiments and revised the manuscript. K.H.M.S, carried out experiments and performed bioinformatic analysis; M.S.V., performed bioinformatic analysis; S.J.K., carried out experiments; R.A., provided technical assistance; G.G., designed and performed experiments, supervised parts of the study and revised manuscript; D.J.A., and B.J.P.H., supervised parts of the study and revised manuscript; F.J.C.-N. designed and performed experiments, supervised the study and revised the manuscript; N.K.W., designed, performed and supervised experiments, and wrote the manuscript; B.G. designed and supervised experiments and wrote the manuscript.

## Corresponding authors

Further information and requests should be directed to and will be fulfilled by the corresponding authors, Nicola K Wilson and Berthold Gottgens.

## Acknowledgements

The authors thank Reiner Schulte, Chiara Cossetti and Gabriela Grondys-Kotarba from the Cambridge Institute for Medical Research Flow Cytometry Core facility for their assistance with cell sorting. We would also like to thank Cambridge Stem Cell Institute Genomics Facility for generating the 10x Genomics libraries and CRUK CI Genomics core for performing high-throughput sequencing, as well as the Cambridge Stem Cell Institute Histology Core for processing of histological samples. Work in the Göttgens laboratory is supported by Wellcome Trust (206328/Z/17/Z), Blood Cancer UK (18002), Cancer Research UK (C1163/A21762). Marija Zaroscinceva was supported by CRUK Cambridge Cancer Centre PhD fellowship (C9685/A29288). Moosa Qureshi was supported by CRUK Cambridge Cancer Centre PhD fellowship (C48525/A18345). Katherine HM Sturgess was supported by Wellcome Trust Clinical PhD Fellowship (204017/Z/16/Z).

Funding in the laboratory of B.J.P.H. was supported by the Leukemia & Lymphoma Society (7035-24), the NIHR Cambridge Biomedical Research Centre (BRC-1215-20014) and the Cancer Research UK Cambridge Centre (CTRQQR-2021\100012). The views expressed are those of the authors and not necessarily those of the NIHR or the Department of Health and Social Care.

This research was funded in whole, or in part, by the Wellcome Trust [203151/Z/16/Z, 203151/A/16/Z] and the UKRI Medical Research Council [MC_PC_17230]. For the purpose of open access, the author has applied a CC BY public copyright licence to any Author Accepted Manuscript version arising from this submission.

## References

1. Williams, N., Lee, J., Mitchell, E., Moore, L., Baxter, E.J., Hewinson, J., Dawson, K.J., Menzies, A., Godfrey, A.L., Green, A.R., et al. (2022). Life histories of myeloproliferative neoplasms inferred from phylogenies. Nature 602. 10.1038/s41586-021-04312-6.

2. Sousos, N., Ní Leathlobhair, M., Simoglou Karali, C., Louka, E., Bienz, N., Royston, D., Clark, S.A., Hamblin, A., Howard, K., Mathews, V., et al. (2022). In utero origin of myelofibrosis presenting in adult monozygotic twins. Nat Med 28. 10.1038/s41591-022-01793-4.

3. Fialkow, P.J., Singer, J.W., Raskind, W.H., Adamson, J.W., Jacobson, R.J., Bernstein, I.D., Dow, L.W., Najfeld, V., and Veith, R. (1987). Clonal Development, Stem-Cell Differentiation, and Clinical Remissions in Acute Nonlymphocytic Leukemia. New England Journal of Medicine 317. 10.1056/nejm198708203170802.

4. Ding, L., Ley, T.J., Larson, D.E., Miller, C.A., Koboldt, D.C., Welch, J.S., Ritchey, J.K., Young, M.A., Lamprecht, T., McLellan, M.D., et al. (2012). Clonal evolution in relapsed acute myeloid leukaemia revealed by whole-genome sequencing. Nature 481. 10.1038/nature10738.

5. Ford, A.M., Mansur, M.B., Furness, C.L., Van Delft, F.W., Okamura, J., Suzuki, T., Kobayashi, H., Kaneko, Y., and Greaves, M. (2015). Protracted dormancy of pre-leukemic stem cells. Leukemia 29. 10.1038/leu.2015.132.

6. Corces-Zimmerman, M.R., Hong, W.J., Weissman, I.L., Medeiros, B.C., and Majeti, R. (2014). Preleukemic mutations in human acute myeloid leukemia affect epigenetic regulators and persist in remission. Proc Natl Acad Sci 111. 10.1073/pnas.1324297111.

7. Shlush, L.I., Zandi, S., Mitchell, A., Chen, W.C., Brandwein, J.M., Gupta, V., Kennedy, J.A., Schimmer, A.D., Schuh, A.C., Yee, K.W., et al. (2014). Identification of pre-leukaemic haematopoietic stem cells in acute leukaemia. Nature 506. 10.1038/nature13038.

8. Lavau, C., Szilvassy, S.J., Slany, R., and Cleary, M.L. (1997). Immortalization and leukemic transformation of a myelomonocytic precursor by retrovirally transduced HRX-ENL. EMBO Journal 16. 10.1093/emboj/16.14.4226.

9. Huntly, B.J.P., Shigematsu, H., Deguchi, K., Lee, B.H., Mizuno, S., Duclos, N., Rowan, R., Amaral, S., Curley, D., Williams, I.R., et al. (2004). MOZ-TIF2, but not BCR-ABL, confers properties of leukemic stem cells to committed murine hematopoietic progenitors. Cancer Cell 6. 10.1016/j.ccr.2004.10.015.

10. Redecke, V., Wu, R., Zhou, J., Finkelstein, D., Chaturvedi, V., High, A.A., and Häcker, H. (2013). Hematopoietic progenitor cell lines with myeloid and lymphoid potential. Nat Methods 10. 10.1038/nmeth.2510.

11. Goardon, N., Marchi, E., Atzberger, A., Quek, L., Schuh, A., Soneji, S., Woll, P., Mead, A., Alford, K.A., Rout, R., et al. (2011). Coexistence of LMPP-like and GMP-like leukemia stem cells in acute myeloid leukemia. Cancer Cell 19. 10.1016/j.ccr.2010.12.012.

12. Basilico, S., Wang, X., Kennedy, A., Tzelepis, K., Giotopoulos, G., Kinston, S.J., Quiros, P.M., Wong, K., Adams, D.J., Carnevalli, L.S., et al. (2020). Dissecting the early steps of MLL induced leukaemogenic transformation using a mouse model of AML. Nat Commun 11. 10.1038/s41467-020-15220-0.

13. Braun, T.P., Okhovat, M., Coblentz, C., Carratt, S.A., Foley, A., Schonrock, Z., Smith, B.M., Nevonen, K., Davis, B., Garcia, B., et al. (2019). Myeloid lineage enhancers drive oncogene synergy in CEBPA/CSF3R mutant acute myeloid leukemia. Nat Commun 10. 10.1038/s41467-019-13364-2.

14. Lin, L.I., Chen, C.Y., Lin, D.T., Tsay, W., Tang, J.L., Yeh, Y.C., Shen, H.L., Su, F.H., Yao, M., Huang, S.Y., et al. (2005). Characterization of CEBPA mutations in acute myeloid leukemia: Most patients with CEBPA mutations have biallelic mutations and show a distinct immunophenotype of the leukemic cells. Clinical Cancer Research 11. 10.1158/1078-0432.CCR-04-1816.

15. Togami, K., Kitaura, J., Uchida, T., Inoue, D., Nishimura, K., Kawabata, K.C., Nagase, R., Horikawa, S., Izawa, K., Fukuyama, T., et al. (2015). A C-terminal mutant of CCAAT-enhancer-binding protein α (C/EBPα-Cm) downregulates Csf1r, a potent accelerator in the progression of acute myeloid leukemia with C/EBPα-Cm. Exp Hematol 43. 10.1016/j.exphem.2014.11.011.

16. Prior, I.A., Lewis, P.D., and Mattos, C. (2012). A comprehensive survey of ras mutations in cancer. Cancer Res 72. 10.1158/0008-5472.CAN-11-2612.

17. Prior, I.A., Hood, F.E., and Hartley, J.L. (2020). The frequency of ras mutations in cancer. Cancer Res 80. 10.1158/0008-5472.CAN-19-3682.

18. Tyner, J.W., Erickson, H., Deininger, M.W.N., Willis, S.G., Eide, C.A., Levine, R.L., Heinrich, M.C., Gattermann, N., Gilliland, D.G., Druker, B.J., et al. (2009). High-throughput sequencing screen reveals novel, transforming RAS mutations in myeloid leukemia patients. Blood 113. 10.1182/blood-2008-04-152157.

19. Stasik, S., Eckardt, J.N., Kramer, M., Röllig, C., Krämer, A., Scholl, S., Hochhaus, A., Crysandt, M., Brümmendorf, T.H., Naumann, R., et al. (2021). Impact of PTPN11 mutations on clinical outcome analyzed in 1529 patients with acute myeloid leukemia. Blood Adv 5. 10.1182/BLOODADVANCES.2021004631.

20. Tartaglia, M., Martinelli, S., Cazzaniga, G., Cordeddu, V., Iavarone, I., Spinelli, M., Palmi, C., Carta, C., Pession, A., Aricò, M., et al. (2004). Genetic evidence for lineage-related and differentiation stage-related contribution of somatic PTPN11 mutations to leukemogenesis in childhood acute leukemia. Blood 104. 10.1182/blood-2003-11-3876.

21. Kratz, C.P., Niemeyer, C.M., Castleberry, R.P., Cetin, M., Bergsträsser, E., Emanuel, P.D., Hasle, H., Kardos, G., Klein, C., Kojima, S., et al. (2005). The mutational spectrum of PTPN11 in juvenile myelomonocytic leukemia and Noonan syndrome/myeloproliferative disease. Blood 106. 10.1182/blood-2005-02-0531.

22. Tartaglia, M., Niemeyer, C.M., Fragale, A., Song, X., Buechner, J., Jung, A., Hählen, K., Hasle, H., Licht, J.D., and Gelb, B.D. (2003). Somatic mutations in PTPN11 in juvenile myelomonocytic leukemia, myelodysplastic syndromes and acute myeloid leukemia. Nat Genet 34. 10.1038/ng1156.

23. Muñoz-Maldonado, C., Zimmer, Y., and Medová, M. (2019). A comparative analysis of individual ras mutations in cancer biology. Front Oncol 9. 10.3389/fonc.2019.01088.

24. Alfayez, M., Issa, G.C., Patel, K.P., Wang, F., Wang, X., Short, N.J., Cortes, J.E., Kadia, T., Ravandi, F., Pierce, S., et al. (2021). The Clinical impact of PTPN11 mutations in adults with acute myeloid leukemia. Leukemia 35. 10.1038/s41375-020-0920-z.

25. Matsuda, K., Shimada, A., Yoshida, N., Ogawa, A., Watanabe, A., Yajima, S., Iizuka, S., Koike, K., Yanai, F., Kawasaki, K., et al. (2007). Spontaneous improvement of hematologic abnormalities in patients having juvenile myelomonocytic leukemia with specific RAS mutations. Blood 109. 10.1182/blood-2006-09-046649.

26. Kato, M., Yasui, N., Seki, M., Kishimoto, H., Sato-Otsubo, A., Hasegawa, D., Kiyokawa, N., Hanada, R., Ogawa, S., Manabe, A., et al. (2013). Aggressive transformation of juvenile myelomonocytic leukemia associated with duplication of oncogenic KRAS due to acquired uniparental disomy. Journal of Pediatrics 162. 10.1016/j.jpeds.2013.01.003.

27. Flotho, C., Kratz, C.P., Bergsträsser, E., Hasle, H., Starý, J., Trebo, M., Van Den Heuvel-Eibrink, M.M., Wójcik, D., Zecca, M., Locatelli, F., et al. (2008). Genotype-phenotype correlation in cases of juvenile myelomonocytic leukemia with clonal RAS mutations. Blood 111. 10.1182/blood-2007-09-111831.

28. Wintering, A., Dvorak, C.C., Stieglitz, E., and Loh, M.L. (2021). Juvenile myelomonocytic leukemia in the molecular era: A clinician’s guide to diagnosis, risk stratification, and treatment. Blood Adv 5. 10.1182/bloodadvances.2021005117.

29. Chao, A.K., Meyer, J.A., Lee, A.G., Hecht, A., Tarver, T., Van Ziffle, J., Koegel, A.K., Golden, C., Braun, B.S., Sweet-Cordero, E.A., et al. (2020). Fusion driven JMML: a novel CCDC88C–FLT3 fusion responsive to sorafenib identified by RNA sequencing. Leukemia 34. 10.1038/s41375-019-0549-y.

30. Gratias, E.J., Liu, Y.L., Meleth, S., Castleberry, R.P., and Emanuel, P.D. (2005). Activating FLT3 mutations are rare in children with juvenile myelomonocytic leukemia. Pediatr Blood Cancer 44. 10.1002/pbc.20176.

31. Nabarro, S., Thrasher, A.J., Kempski, H., Amrolia, P., and Anderson, J. (2003). Generation of immunostimulatory dendritic cells from the malignant clone in patients with juvenile myelomonocytic leukemia. Leukemia 17. 10.1038/sj.leu.2403059.

32. Longoni, D., D’Amico, G., Gaipa, G., Bernasconi, S., Vulcano, M., Onnis, P., Niemeyer, C.M., Allavena, P., and Biondi, A. (2002). Commitment of juvenile myelo-monocytic (JMML) leukemic cells to spontaneously differentiate into dendritic cells. Hematology Journal 3. 10.1038/sj.thj.6200192.

33. Meyerson, H., Awadallah, A., Blidaru, G., Osei, E., Schlegelmilch, J., Egler, R., Abu-Arja, R., and Ding, H. (2017). Juvenile myelomonocytic leukemia with prominent CD141+ myeloid dendritic cell differentiation. Hum Pathol 68. 10.1016/j.humpath.2017.03.025.

34. Chan, R.J., Cooper, T., Kratz, C.P., Weiss, B., and Loh, M.L. (2009). Juvenile myelomonocytic leukemia: A report from the 2nd International JMML Symposium. Leuk Res 33. 10.1016/j.leukres.2008.08.022.

35. Niemeyer, C.M. (2018). JMML genomics and decisions. Hematology (United States) 2018. 10.1182/asheducation-2018.1.307.

36. Caye, A., Rouault-Pierre, K., Strullu, M., Lainey, E., Abarrategi, A., Fenneteau, O., Arfeuille, C., Osman, J., Cassinat, B., Pereira, S., et al. (2020). Despite mutation acquisition in hematopoietic stem cells, JMML-propagating cells are not always restricted to this compartment. Leukemia 34. 10.1038/s41375-019-0662-y.

37. Hartmann, M., Schönung, M., Rajak, J., Maurer, V., Hai, L., Bauer, K., Hakobyan, M., Staeble, S., Langstein, J., Jardine, L., et al. (2023). Oncogenic RAS-Pathway Activation Drives Oncofetal Reprogramming and Creates Therapeutic Vulnerabilities in Juvenile Myelomonocytic Leukemia. bioRxiv, 2023.10.27.563754. 10.1101/2023.10.27.563754.

38. Hashmi, S.K., Punia, J.N., Gaikwad, A.S., Fisher, K.E., Roy, A., Lopez-Terrada, D.H., Rao, P., Ringrose, J., Marcogliese, A.N., and Rau, R.E. (2016). Aberrant Precursor B Lymphoid Blast Population in a Patient with Juvenile Myelomonocytic Leukemia. Blood 128. 10.1182/blood.v128.22.5557.5557.

39. Lau, R.C., Squire, J., Brisson, L., Kamel-Reid, S., Grunberger, T., Dubé, I., Letarte, M., Shannon, K., and Freedman, M.H. (1994). Lymphoid blast crisis of B-lineage phenotype with monosomy 7 in a patient with juvenile chronic myelogenous leukemia (JCML). Leukemia 8.

40. Zeissig, M.N., Ashwood, L.M., Kondrashova, O., and Sutherland, K.D. (2023). Next batter up! Targeting cancers with KRAS-G12D mutations. Trends Cancer 9. 10.1016/j.trecan.2023.07.010.

41. Huang, L., Guo, Z., Wang, F., and Fu, L. (2021). KRAS mutation: from undruggable to druggable in cancer. Signal Transduct Target Ther 6. 10.1038/s41392-021-00780-4.

42. Sakashita, K., Kato, I., Daifu, T., Saida, S., Hiramatsu, H., Nishinaka, Y., Ebihara, Y., Ma, F., Matsuda, K., Saito, S., et al. (2015). In vitro expansion of CD34+ CD38-cells under stimulation with hematopoietic growth factors on AGM-S3 cells in juvenile myelomonocytic leukemia. Leukemia 29. 10.1038/leu.2014.239.

43. Picelli, S., Faridani, O.R., Björklund, Å.K., Winberg, G., Sagasser, S., and Sandberg, R. (2014). Full-length RNA-seq from single cells using Smart-seq2. Nat Protoc 9. 10.1038/nprot.2014.006.

44. Bagnoli, J.W., Ziegenhain, C., Janjic, A., Wange, L.E., Vieth, B., Parekh, S., Geuder, J., Hellmann, I., and Enard, W. (2018). Sensitive and powerful single-cell RNA sequencing using mcSCRB-seq. Nat Commun 9. 10.1038/s41467-018-05347-6.

45. Kucinski, I., Wilson, N.K., Hannah, R., Kinston, S.J., Cauchy, P., Lenaerts, A., Grosschedl, R., and Göttgens, B. (2020). Interactions between lineage-associated transcription factors govern haematopoietic progenitor states. EMBO J 39. 10.15252/embj.2020104983.

46. Wolf, F.A., Angerer, P., and Theis, F.J. (2018). SCANPY: Large-scale single-cell gene expression data analysis. Genome Biol 19. 10.1186/s13059-017-1382-0.

47. Kucinski, I., Campos, J., Barile, M., Severi, F., Bohin, N., Moreira, P.N., Allen, L., Lawson, H., Haltalli, M.L.R., Kinston, S.J., et al. (2024). A time- and single-cell-resolved model of murine bone marrow hematopoiesis. Cell Stem Cell 31. 10.1016/j.stem.2023.12.001.

48. Wolock, S.L., Lopez, R., and Klein, A.M. (2019). Scrublet: Computational Identification of Cell Doublets in Single-Cell Transcriptomic Data. Cell Syst 8. 10.1016/j.cels.2018.11.005.

49. Stuart, T., Butler, A., Hoffman, P., Hafemeister, C., Papalexi, E., Mauck, W.M., Hao, Y., Stoeckius, M., Smibert, P., and Satija, R. (2019). Comprehensive Integration of Single-Cell Data. Cell 177. 10.1016/j.cell.2019.05.031.

50. McInnes, L., Healy, J., and Melville, J. (2020). UMAP: Uniform Manifold Approximation and Projection for Dimension Reduction. ArXiv.

51. Li, H. (2013). Aligning sequence reads, clone sequences and assembly contigs with BWA-MEM.

52. Auwera, G. van der, and O’Connor, B.D. (2020). Genomics in the cloud : using Docker, GATK, and WDL in Terra First edition. (O’Reilly Media).

53. Danecek, P., Bonfield, J.K., Liddle, J., Marshall, J., Ohan, V., Pollard, M.O., Whitwham, A., Keane, T., McCarthy, S.A., and Davies, R.M. (2021). Twelve years of SAMtools and BCFtools. Gigascience 10. 10.1093/gigascience/giab008.

54. Wang, K., Li, M., and Hakonarson, H. (2010). ANNOVAR: Functional annotation of genetic variants from high-throughput sequencing data. Nucleic Acids Res 38. 10.1093/nar/gkq603.

55. Mayakonda, A., Lin, D.C., Assenov, Y., Plass, C., and Koeffler, H.P. (2018). Maftools: Efficient and comprehensive analysis of somatic variants in cancer. Genome Res 28. 10.1101/gr.239244.118.

56. Love, M.I., Huber, W., and Anders, S. (2014). Moderated estimation of fold change and dispersion for RNA-seq data with DESeq2. Genome Biol 15. 10.1186/s13059-014-0550-8.

